# Personalized composite scaffolds for accelerated cell- and growth factor-free craniofacial bone regeneration

**DOI:** 10.1101/2024.02.18.580898

**Authors:** Mirae Kim, Caralyn P. Collins, Yugang Liu, Hsiu-Ming Tsal, Yujin Ahn, Xinlong Wang, Joseph W. Song, Chongwen Duan, Cheng Sun, Zhu Yi, Tong-Chuan He, Russell R. Reid, Guillermo A. Ameer

## Abstract

Approaches to regenerating bone often rely on the integration of biomaterials and biological signals in the form of cells or cytokines. However, from a translational point of view, these approaches face challenges due to the sourcing and quality of the biologic, unpredictable immune responses, complex regulatory paths, and high costs. We describe a simple manufacturing process and a material-centric 3D-printed composite scaffold system (CSS) that offers distinct advantages for clinical translation. The CSS comprises a 3D-printed porous polydiolcitrate-hydroxyapatite composite elastomer infused with a polydiolcitrate-graphene oxide hydrogel composite. Using a continuous liquid interface production 3D printer, we fabricate a precise porous ceramic scaffold with 60% hydroxyapatite content resembling natural bone. The resulting scaffold integrates with a thermoresponsive hydrogel composite, customizable *in situ* to fit the defect. This hybrid phasic porous CSS mimics the bone microenvironment (inorganic and organic) while allowing independent control of each material phase (rigid and soft). The CSS stimulates osteogenic differentiation *in vitro* and *in vivo*. Moreover, it promotes M2 polarization and blood vessel ingrowth, which are crucial for supporting bone formation. Our comprehensive micro-CT analysis revealed that within 4 weeks in a critical-size defect model, the CSS accelerated ECM deposition (8-fold) and mineralized osteoid (69-fold) compared to the untreated. Our material-centric approach delivers impressive osteogenic properties and streamlined manufacturing advantages, potentially expediting clinical application for bone reconstruction surgeries.

## Introduction

Over 3 million cases of craniofacial trauma occur each year in the United States, constituting 21% of significant traumas (*1*). Craniofacial bone defects resulting from traumatic injuries present challenges for patients and surgeons, necessitating complex surgeries and substantial surgical costs (*2*). Autografts, while considered the gold standard for craniofacial reconstruction (*3, 4*), bring challenges such as the need for a second/donor surgical site, prolonged operation time, and increased patient discomfort and recovery. Bone tissue engineering has been considered a promising alternative, aiming to replicate the native craniofacial environment without complex technical demands (*5*). However, integrating biological components such as stem cells and growth factors increases regulatory complexity (*6*), raises product development costs, and can elicit unwanted immune and inflammatory responses (*7, 8*). In this regard, material-centric approaches alongside the development of synthetic biological scaffolds offer significant potential for accelerating commercialization strategies and improving patient outcomes (*5, 9*).

Bone extracellular matrix (ECM) is composed of approximately 40% organic and 60% inorganic compounds (*10*). Several material-centric strategies have proposed composite scaffold systems (**CSS**) to accelerate osteogenesis and vascularization, either by integrating ceramics with hydrogels to simulate the bone matrix microenvironment (*11*) or by combining them with polymers and additives (e.g., bioactive particles or graphene derivates) (*12, 13*). However, the low solubility of ceramics under physiological conditions leads to poor interface interaction with hydrogels, causing structural instability, while employing particles for enhancing miscibility (*14, 15*) might compromise biocompatibility. Pre-fabricated systems incorporating hydrogels should consider swelling properties (*16, 17*), which may lead to structural misalignment post-implantation. Moreover, the mechanical properties of porous ceramic scaffolds can be influenced by the quantity of additives (*12, 18*), complicating the independent control of structural and functional properties.

CSS should not only provide a microenvironment conducive to osteogenesis but also conform to the defect geometry in 3D for effective integration and function recovery (*19*). Additive manufacturing using continuous liquid interface production (CLIP) offers advantages in printing speed and complex architecture fabrication at high resolutions (*20, 21*). Therefore, 3D-printed CSS developed using CLIP technology has significant potential in bone reconstruction. Nevertheless, several complex considerations are required for this manufacturing process, including additional steps to improve material compatibility among the composite components (*14*). Due to the aforementioned reasons, material-centric strategies involving composites face numerous considerations and constraints.

Herein, we present a customizable CSS comprised of biphasic citrate-based polymers, incorporating two microparticles, hydroxyapatite (**HA**) and graphene oxide (**GO**). This CSS is developed by integrating a polydiolcitrate-GO hydrogel composite into a 3D-printed porous polydiolcitrate-HA scaffold. The 3D-printed scaffold utilizes a poly(1,8-octanediol citrate) (**POC**) citrate elastomer, which exhibits superior cytocompatibility and tissue interaction compared to commonly used poly(caprolactone) (PCL) in bone tissue engineering (*22*). The polydiolcirate-GO hydrogel composite, a thermal-responsive gel, provides shaping flexibility, enabling seamless integration with the 3D-printed scaffold and *in situ* fabrication to fit the defect site (*23, 24*). We show that the hydrogel-infused 3D-printed CSS stimulates angiogenesis and osteogenesis of endogenous progenitors. Moreover, through granular micro-CT analysis, bone formation over time is assessed in a critical-size defect model in rodents. The material design and strategy allow independent control of each material phase of the CSS and facilitate patient translation and scalable manufacturing, indicating potential as an advanced CSS, particularly for addressing cranial defects.

## Results

### GP hydrogel integrates into P-HA enabling CSS fabrication

The 3D-printed porous ceramic scaffolds are precisely structured to fit a mouse skull defect model via a micro-continuous liquid interface production (μCLIP) 3D printer (**Fig. 1A**). Within this system, the composite mixture is placed in a resin bath, allowing photopolymerization by UV light penetration through an O2-permeable window. Subsequently, the composite is additively manufactured by pattering UV light into cross-sectional images of the 3D-designed scaffold via a digital micromirror device (DMD). Following the printing, the hybrid CSS is fabricated by injecting the hydrogel-GO precursor solution into the structure and gelling it at 37°C. The combination of 3D printing technology (*20, 21*) and the injectable thermoresponsive hydrogel (*23, 25*) allows for the convenient and swift fabrication of a customizable soft-rigid hybrid system.

**Fig. 1.**
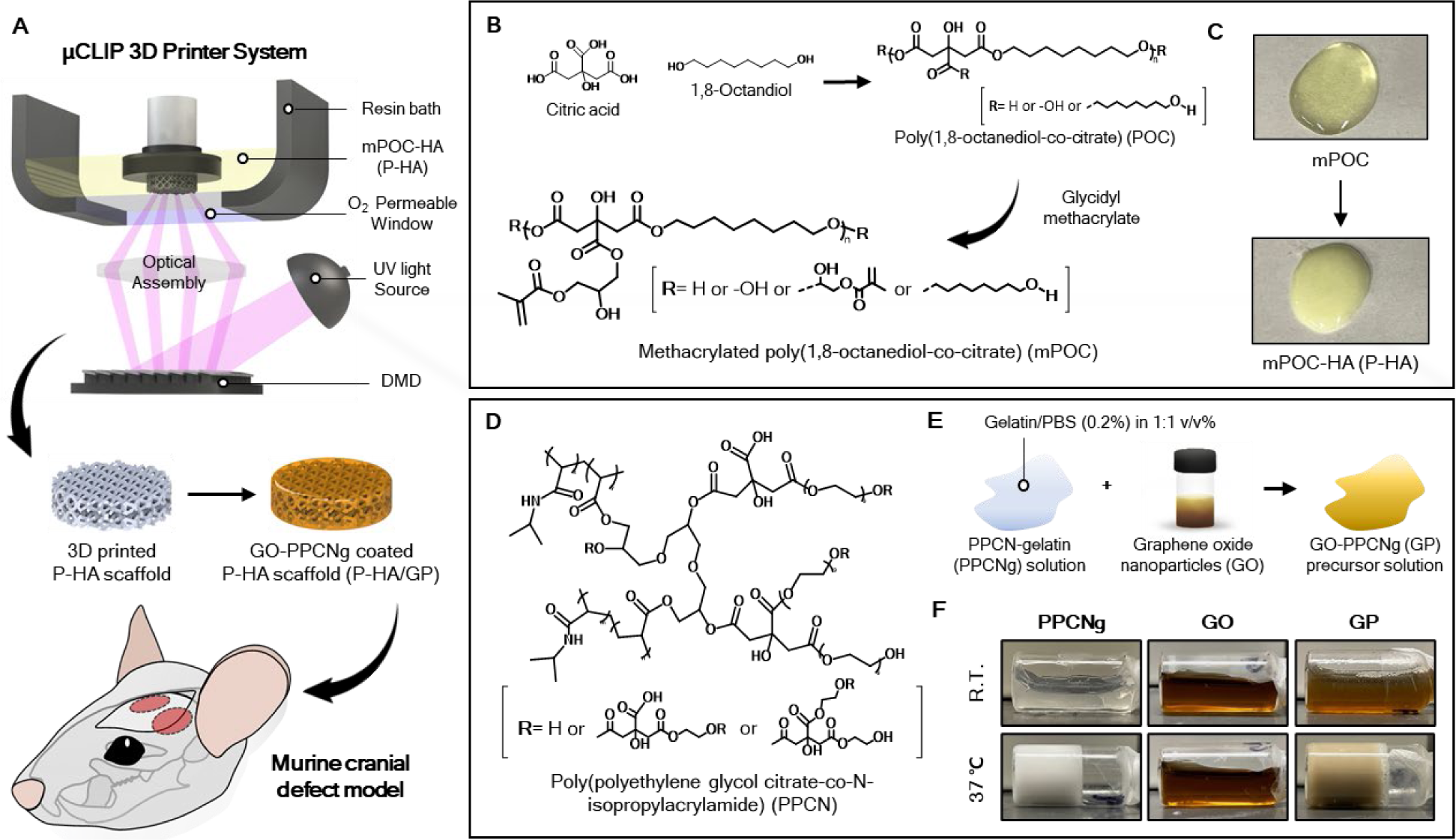
Representative images of the 3D-printed CSS fabrication and preparation of P-HA and GP hydrogel. (A) μCLIP 3D printer system and schematic depiction of 3D-printed CSS implantation in cranial defect model. (B) Synthesis schematic of mPOC and its structure. (C) Preparation of P-HA composite consisting of mPOC polymer and HA. (D) Structure of PPCN and (E) schematic illustration of GP precursor solution preparation. (F) Different physiological properties of GO (0.4mg/mL), PPCNg (50mg/mL), and GP hybrid hydrogel at room temperature and 37℃.

The methacrylated poly(1,8-octanediol-citrate) (**mPOC**) was used as a 3D printable polymer and developed from POC via two steps (**Fig. 1B**). Initially, the POC was obtained through an esterification reaction between 1,8-octanediol and citric acid (*26*). Subsequently, it was developed into mPOC by introducing methacrylate functional groups through a ring-opening reaction of glycidyl methacrylate. The chemical composition of mPOC was confirmed through ^1^H- NMR and FT-IR spectroscopy analysis (**Fig. S1**). The ^1^H-NMR analysis identified peaks at 1.9, 5.7, and 6ppm, indicating the presence of methacrylate groups within the mPOC structure. The estimated molar ratio of citric acid to methacrylate was approximately 1:0.9, as inferred from the spectrum. In addition, the FT-IR spectrum exhibited a C=C stretching vibration peak at 1636 cm^-1^. These results collectively demonstrate the effective modification of POC through methacrylation, allowing for radical polymerization under UV light throughout the 3D printing process.

The formulation of the composite involved a mixture of mPOC with HA microparticles (**P-HA**) (**Fig. 1C**). In order to achieve a wide range of 3D printable composite, we incorporated 2.5μm HA (specific surface area, ≥80 m^2^/g). A previous study has demonstrated that materials derived from HA exhibit cytocompatibility with stem cells and promote the osteogenic differentiation of such cells (*27*). The incorporation of HA microparticles improved the viscosity of the composite, thereby enhancing its printability and allowing for HA contents of up to 60%.

Poly(polyethylene glycol citrate-co-N-isopropylacrylamide) (**PPCN**) was used as the thermoseponsive hydrogel component (**Fig. 1D**). It was obtained with citric acid, PEG and N-isopropylacrylamide (**NIPAAm**) components (*23*), and due to the unique properties of the NIPAAm, it exhibits a lower critical solution temperature (LCST) that enables phase change from liquid to gel at physiological body temperature (37°C) (*28*). The ^1^H-NMR spectrum of PPCN exhibited multiple peaks associated with citric acid, PEG, and NIPAAm units (**Fig. S2**). As a result of analyzing the signal intensity, the molar ratio between citric acid and poly(NIPAAm) was determined to be approximately 1:12, reflecting the molar feed ratio during synthesis. The FT-IR spectrum revealed that PPCN had characteristic amide peaks present in the NIPAAm structure, along with additional peaks indicating C=O, C-O, and -OH functional groups, attributed to citric acid, ester bonds and PEG (**Fig. S2**).

The PPCN was combined with gelatin (PPCNg) and then developed into GO-PPCNg (**GP**) hydrogel composite using the following method. In brief, the PPCN was dissolved in PBS and blended with gelatin in a 1:1 ratio. Afterward, this mixture was combined with GO solution at a volume ratio of 5:1 (**Fig. 1E**). The physical appearance of GP hydrogel demonstrated favorable mixing of GO with PPCNg (**Fig. 1F**). It exhibited the LCST behavior typical of PPCN, remaining in a liquid state at 4°C but undergoing a relatively rapid gelation process at 37°C, while GO solution consistently remained in a liquid form.

### HA and GO improve mechanical and rheological properties of the scaffold

The combination of mPOC and HA provides processing flexibility, rendering it an ideal composite for fabricating 3D-printed porous structures of various dimensions and pore unit cell configurations using the μCLIP 3D printer (**Fig. S3**). For the subsequent experiments, the P-HA was engineered with a porous architecture featuring hexagonal unit cells (**Fig. 2A and Fig. S4**).

**Fig. 2.**
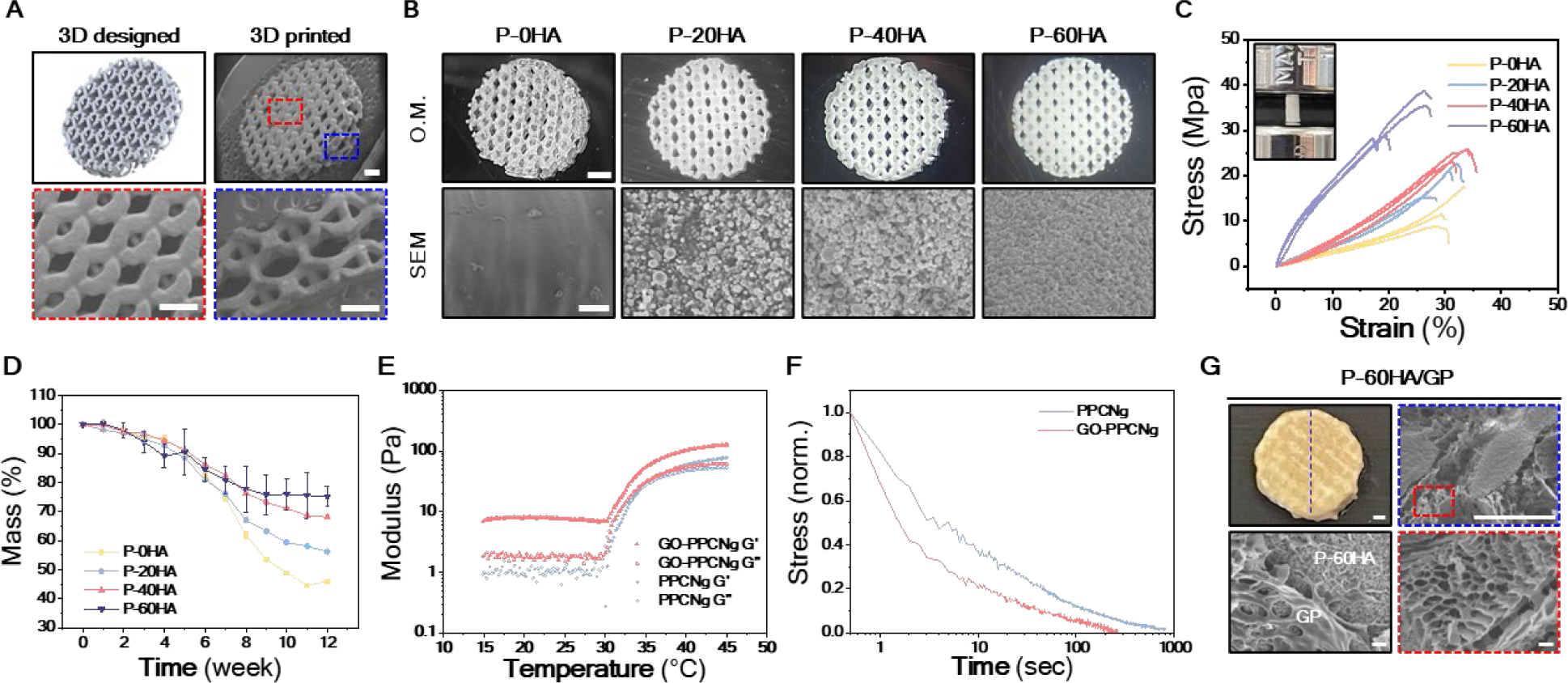
Fabrication of 3D-printed porous P-HA scaffolds and characterization of P-HA scaffolds and GP hydrogel. (A) Comparison images between 3D design and 3D-printed P-HA scaffold. Scale bars, 500μm. (B) 3D-printed P-HA scaffolds with various HA concentrations and morphological surface appearance of each scaffold in SEM. Scale bar, 1μm (top) and 10μm (bottom). (C) Representative stress-strain curves of each P-HA scaffold. n=3 (D) Degradation behavior of each P-HA scaffold at 75°C. Error bars, ± SD; n=3. (E) Gelation kinetics of PPCNg and GP hydrogel, and (F) their stress-relaxation profiles at 37°C. (G) Morphological structure of P-60HA/GP composite scaffold and GP hydrogel in SEM images. Scale bars, 500μm (top) and 10μm (bottom).

The hexagonal structural element efficiently disperses external forces, enhancing the stability of the scaffolds (*29, 30*). Additionally, smaller internal pore units (<450μm) (**Fig. S4**) aimed to promote osteogenesis and vascularization (*31, 32*) while enabling the advantageous integration of the GP hydrogel throughout the scaffold. The μCLIP printer enabled precise customization by replicating the scaffold at a high resolution according to the designed structure without any pore blockages (**Fig. 2A**). The P-HA were prepared at various HA concentrations ranging from 0% to 60% to evaluate mechanical properties according to HA content, and were labeled as P-0HA, P-20H, P-40H, and P-60HA depending on the content. Even with high HA content (60wt.%), the P-HA exhibited favorable printability characteristics (**Fig. S3**), and HA particles were distributed throughout the structure, as depicted in SEM (**Fig. 2B**). The surface roughness resulting from the addition of HA can promote cell adhesion and differentiation (*33, 34*). Therefore, we expected that the surface properties of the P-HA scaffold would provide an advantage in osteogenesis.

In order to assess mechanical properties, P-HA were made into plug-shaped samples (3mm×6mm), which is a standard structure (ASTM D695) for measuring the compressive modulus of materials (**Fig. 2C**). The samples were measured using a universal testing machine until they fractured under a compressive load. Despite the potential impact of higher HA content on sample brittleness, the P-60HA exhibited enhancement in compressive strength (23.7 ± 1.6 MPa) (**Fig. S5**). This enhancement confirms the structural stability of the P-HA composite material, signifying compatibility between mPOC and HA (**Fig. 2B**). The degradation behavior of 3D-printed porous P-HA scaffolds (**Fig. 2A**) was investigated in PBS for 12 weeks at 75°C, representing an environment accelerated 16 times compared to body temperature (*35*) (**Fig. 2D**). At 12 weeks, the P-0HA exhibited a mass loss of approximately 54.1%, while that of P-60HA was 24.6%, which delayed the degradation behavior by about 2.3-fold.

We examined the impact of GO on the rheological and viscoelastic properties of the PPCNg hydrogel (**Fig. 2, E and F**). The PPCNg hydrogels showed a phase transition from liquid to gel at 35°C, where G″ and G′ intersected (**Fig. 2E**). While the GP hydrogel did not show a distinct intersection point, it exhibited a transition from liquid to gel above 35°C (**Fig. 1E**). Furthermore, the GP hydrogel displayed gel-like characteristics with higher G′ values even in the liquid phase (below 35°C) and improved the G′ of the PPCNg hydrogel from 77 to 126 Pa following the phase transition. This behavior is attributed to the interactions between GO and PPCNg chains within the GP mixture (*36*). The stress relaxation of the hydrogels was investigated at 37°C, maintaining a constant shear strain of 15%, comparable to the strain applied by cells within a 3D matrix (*37, 38*) (**Fig. 2F**). The GP hydrogel showed a faster half-stress relaxation time (t1/2 ≈1.5 sec) compared to PPCNg hydrogel (t1/2 ≈3 sec), which is likely due to GO particles interfering with the crosslinking of PPCN chains, leading to faster chain relaxation.

The P-60HA scaffold was combined with GP hydrogel, and the resulting hybrid CSS (**P- 60HA/GP**) maintained a stable composite structure at 37°C (**Fig. 2G**). The morphological structure and the distribution of GP hydrogel within the CSS were evaluated via SEM analysis (**Fig. 2G**). The GP hydrogel formed an extensive network by physically interacting with the P-60HA scaffold and uniformly covered the entire structure. Additionally, the GP hydrogel exhibited permeable porous channels supporting blood vessel formation and tissue ingrowth.

We confirmed that both the P-60HA scaffold and GP hydrogel collectively enhance physical properties, and the hybrid CSS demonstrated a consistent and durable structure. In subsequent experiments, the P-60HA scaffold was chosen as the primary structural framework of the CSS based on results, demonstrating improved mechanical properties with comparable mineral concentration to native bone (65-70wt.%).

### P-60HA/GP is cytocompatible and promotes osteogenesis *in vitro*

We investigated the cytotoxicity and the *in vitro* osteogenic potential of the scaffolds using human mesenchymal stromal cells (hMSCs) (**Fig. 3**). To evaluate the influence of HA and GP hydrogel on cellular activity, the P-0HA, P-60HA, and P-60HA/GP scaffolds were examined, and each value was normalized to TCP control group (**Fig. 3, A and B**). The scaffolds were immersed in the TCP cultured with hMSCs for 7 days and subjected to live/dead staining (**Fig. 3A**) and alamarBlue assay **(Fig. 3B**). There were no observable dead cells, with sustained live cell proliferation, and cell viability was recorded at ≈90% for 7 days, suggesting that both P-HA scaffolds and GP hydrogel are biocompatible.

**Fig. 3.**
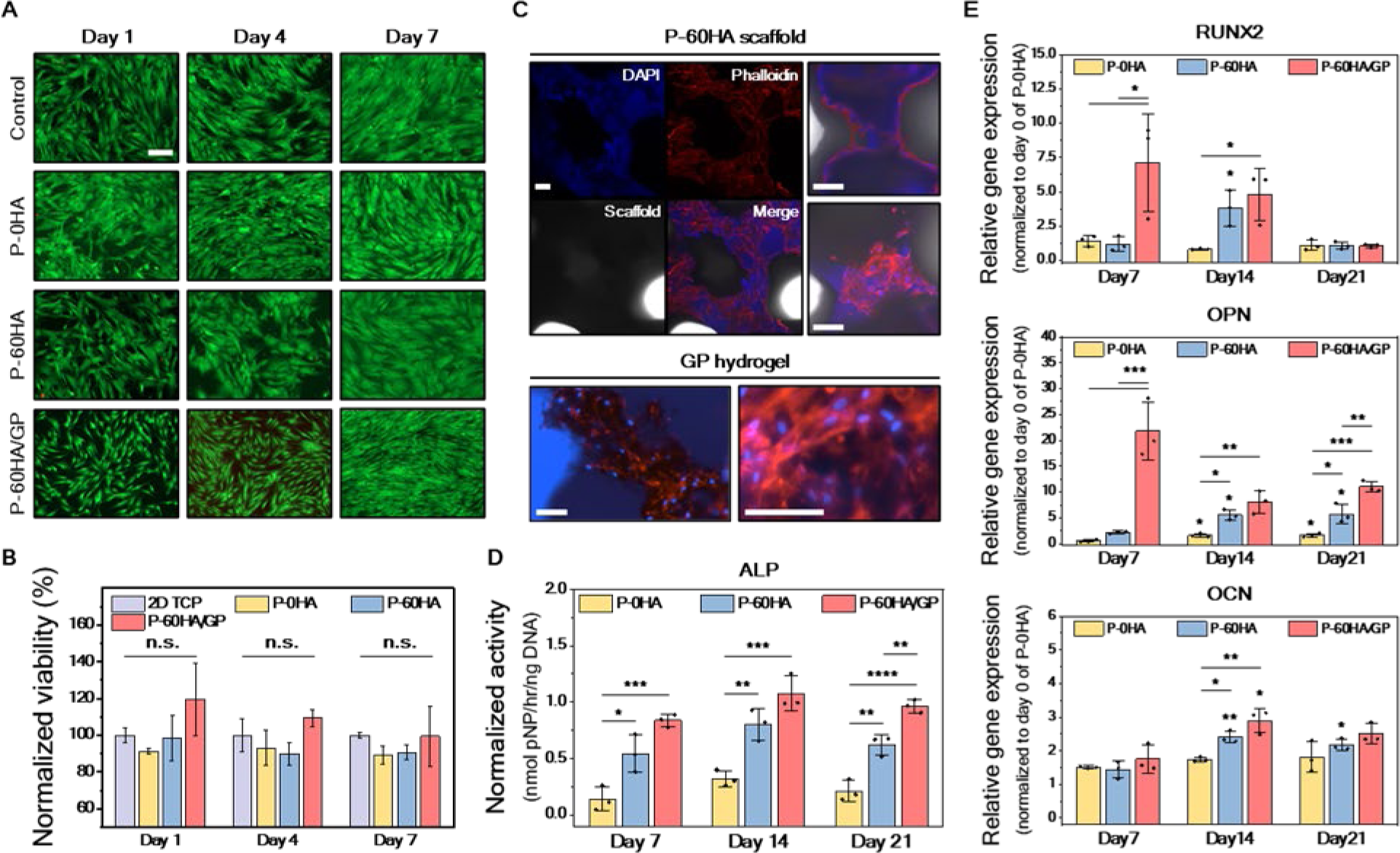
In vitro assessment of cell viability and osteogenic differentiation of hMSCs on the scaffolds. (A) Live/dead staining images of each group on days 1, 4, and 7 days. Scale bar, 200μm. (B) Cell viability in the alamarBlue assay normalized to the TCP control group. n.s.: no significant difference; Error bars, ± SD; n=3. (C) Cytoskeleton staining images of hMSCs on P-60HA scaffold and GP hydrogel in P-60HA/GP scaffold 4 days after cell seeding. Scale bars, 100μm. (D) ALP activity normalized to DNA concentration in each group on days 7, 14, and 21. \**p* <0.05, \*\**p* <0.01, \*\*\**p* <0.001 and \*\*\*\**p* <0.0001; Error bars, ± SD; n=3. (E) The relative expression levels of RUNX2, OPN, and OCN for hMSCs cultured in each group at days 7, 14, and 21. All expression levels were quantified using 2–ΔΔCT method and then normalized to the value of the housekeeping gene GAPDH and day 0 for the P-0HA group. \**p* <0.05, \*\**p* <0.01, \*\*\**p* <0.001 and \*\*\*\**p* <0.0001; Error bars, ± SD; n=3.

The interaction between scaffolds and cells plays a crucial role as it enhances tissue reconstruction by enabling effective interaction with surrounding tissues post-implantation (*39, 40*). We assessed cell adhesion and retention on the scaffolds using cytoskeleton staining (**Fig. 3C**). hMSCs were cultured on P-60HA and P-60HA/GP for 4 days without cell adhesion treatment.

In the staining results, P-60HA exhibited cell attachment, but cells had limited proliferation on the scaffold surface. In contrast, cells showed widespread distribution within the GP hydrogel of P- 60HA/GP. This indicates that GP hydrogel provides a conducive microenvironment to cell growth, allowing cells to proliferate throughout the GP hydrogel-conjugated scaffold.

To demonstrate the effects of HA and GP hydrogel on the osteogenic differentiation of hMSCs, we analyzed osteogenesis markers at 7, 14, and 21 days after cell culturing on the scaffolds. First, the early osteogenic marker alkaline phosphatase (ALP) was assessed using the absorbance method and normalized to the DNA concentration of each group at the indicated time points (**Fig. 3D**). At day 7, ALP activity was significantly upregulated in P-60HA and P-60HA/GP compared to P-0HA. Moreover, the incorporation of GP hydrogel into the P-60HA scaffold enhanced ALP activity, which was ≈4.57 times (\*\*\*\**p* <0.0001) higher than P-0HA at day 21.

The expression levels of osteogenesis markers were analyzed by real-time reverse quantitative PCR (RT-qPCR), and each value was normalized to day 0 of P-0HA scaffold for comparison between groups (**Fig. 3E**). The results showed that P-60HA/GP upregulated early (Runt-related transcription factor 2, RUNX2) (≈5-fold) and intermediate (Osteopontin, OPN) markers (≈39-fold) compared to P-0HA from the early time point, day 7. By day 14, while the trend in OPN levels differed in P-60HA/GP from the other scaffolds, it maintained 5- and 1.5-fold higher levels than those of P-0HA and P-60HA, respectively. The late-stage marker, osteocalcin (OCN), exhibited upregulation at day 14 in both P-60HA and P-60HA/GP, with the P-60HA/GP showing enhanced levels 1.7 times compared to P-0HA.

Taken together, P-60HA/GP induced early osteogenesis with high levels of ALP activity, RUNX2, and OPN during the proliferative phase before mineralization and upregulated OCN promoting apatite mineralization *in vitro* (*41*).

### P-60HA/GP exhibits favorable tissue interaction and biocompatibility *in vivo*

We investigated the immune response and tissue interaction associated with P-HA scaffolds through subcutaneous implantation in a mouse model (**Fig. 4A**). Each scaffold was implanted on the dorsal region of the mice, and tissue samples were harvested with scaffolds after 7 and 35 days for histological analysis. Following *in vivo* implantation, the infiltration of cells within the scaffold serves as an indicator of the scaffold’s capability to facilitate cell attachment, proliferation, and migration within its structure (*42*). H&E staining results (**Fig. 4B**) revealed mild connective tissue and cellular infiltration into the porous scaffold structure by day 7 in all experimental groups, and there were no significant inflammatory responses at the implantation site. Notably, P-60HA/GP exhibited robust cell infiltration at the administered GP hydrogel area. This cell recruitment can be attributed to the favorable effects of the gelatin (*24*) and GO components (*43*) in the GP hydrogel. By day 35, all scaffolds demonstrated successful integration with the surrounding tissue network, and some biomaterial-associated multinucleated giant cells (BMGCs) were observed around the surface of the scaffolds. These cells are typically observed in response to foreign body reactions of polymeric implants (*40, 44*). BMGCs may act as key regulators during biomaterial integration and have the potential to contribute to the vascularization of the implant bed, ultimately stimulating bone formation (*44*). This observation suggests that P-HA scaffolds interact with surrounding tissues and induce cellular responses, contributing to tissue remodeling.

**Fig. 4.**
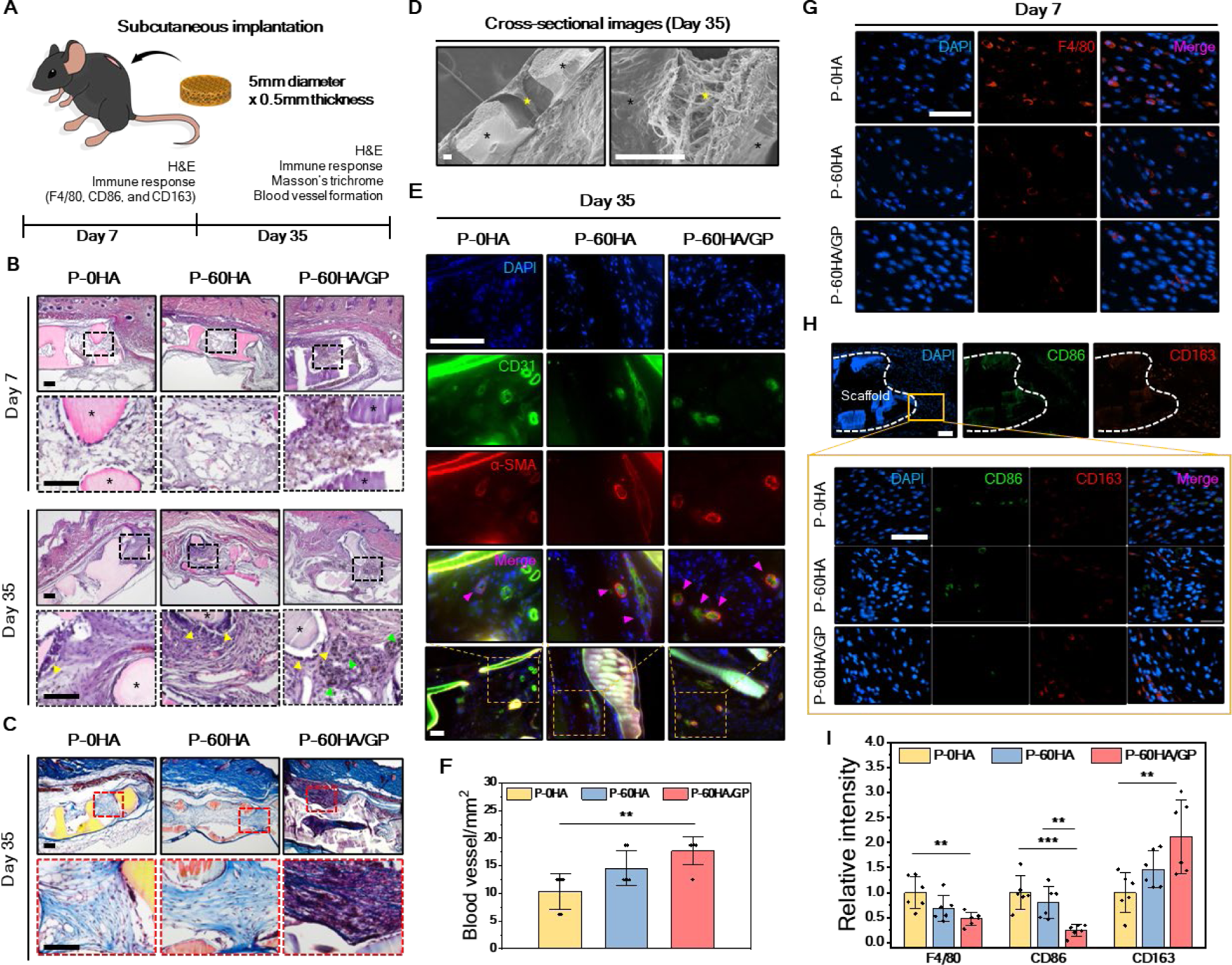
Evaluation of in vivo immune response and biocompatibility of scaffolds in mouse subcutaneous implantation. (A) Schematic illustration of the subcutaneous implantation experiment. (B) Cross-sectional H&E histological images of scaffold implanted tissue at day 7 and day 35. The images at the bottom represent the higher magnification of each group. The asterisk (*): scaffold; Green arrowhead: GO residue; Yellow arrowhead: multinucleated giant cells. Scale bars, 150μm. (C) Representative Masson’s trichrome staining images at day 35 after implantation and their higher magnification images. The asterisk (*), scaffold; Scale bars, 50μm. (D) Cross-sectional SEM images of the implanted scaffold on day 35. The asterisk (*): scaffold; Star: infiltrated tissues. Left: under 100X magnification; Right: 1,000X magnification. Scale bars, 50μm. (E) Representative immunofluorescence staining images of CD31 and α-SMA on day 35. Pink arrowhead: newly formed blood vessel. Scale bar, 100μm. (F) The quantitative analysis of blood vessel formation inside scaffolds on day 35. \*\**p* <0.01, Error bars, ± SD; n=6. (G) Representative immunofluorescence staining images of F4/80, (H) CD86, and CD163 markers on day 7. Scale bars, 50μm. (I) The relative quantitative mean gray value of F4/80, CD86, and CD163 on day 7. \*\**p* <0.01, and \*\*\**p* <0.001, Error bars, ± SD; n=6.

At 35 days, Masson’s trichrome staining (**Fig. 4C**) was performed to assess the capacity to facilitate effective integration with adjacent tissues and act as a substrate for the deposition of new ECM. All P-HA scaffolds exhibited collagen fibril formation throughout their porous structures. Specifically, within the P-60HA/GP scaffold, there was observable tissue infiltration aligned along the site of GP hydrogel injection. This observation is attributed to the interaction between GO and collagen fibers (*45*), such as hydrogen bonding, electrostatic interaction, and π-π stacking (*46, 47*). The stable ECM network formation surrounding the scaffolds and within their structure was also confirmed in SEM (**Fig. 4D**). Overall, these results showed the potential of P- HA scaffolds and GP hydrogel to stimulate cell and tissue ingrowth over time, indicating their suitability as an implant for tissue reconstruction.

### GP hydrogel affects angiogenesis and M2 macrophage polarization

Timely vascularization supplies oxygen and nutrients during bone repair, thereby enhancing bone formation (*48*). To further evaluate their potential in stimulating angiogenesis, we conducted immunofluorescence (IF) analysis on the tissue formed within the porous structure of the scaffolds (n=6 from three biologically independent mice) (**Fig. 4E**). At 35 days post-implantation, there was no statistically significant difference (*p* =0.17) observed between P-0HA (10 ± 3 vessels/mm^2^) and P-60HA (15 ± 3 vessels/mm^2^) as a quantitative result by CD31 and α-smooth muscle actin (α- SMA) markers (**Fig. 4F**). However, P-60HA/GP demonstrated enhanced blood vessel formation (18 ± 3 vessels/mm^2^) (\*\**p* <0.01) compared to the P-0HA, suggesting that the inclusion of GP hydrogel accelerates angiogenesis (*49, 50*).

Once the biomaterial is implanted, an inflammatory reaction is observed for foreign body response (*51*). Macrophages are one of the first cells to encounter the implanted materials and the major modulator of tissue integration (*52*), and they exhibit a wide range of capabilities, capable of transitioning from an M1 type (pro-inflammatory state) to an M2 type (anti-inflammatory state) (*53*). During the bone regeneration period, the long-term M1 macrophage environment after implantation may lead to bone destruction, hindering the process of bone regeneration and repair. We evaluated the degree of inflammation and macrophage polarization using IF staining with F4/80 (pan macrophages), CD86 (M1 macrophages), and CD163 (M2 macrophages) (n=6 from three biologically independent mice) (**Fig. 4, G and H**). The analysis was performed on the adjacent tissues surrounding the implanted scaffold, and the relative mean gray value was determined based on the P-0HA value at day 7 (**Fig. 4I**). At 35 days, F4/80 and CD86 levels were decreased in all groups (mean value <0.5) (**Fig. S6**) by the transition from the pro-inflammatory to anti-inflammatory phase, and there was no significant difference between groups (*p* >0.05). However, on day 7, P-60HA/GP showed low F4/80 intensity (0.5 ± 0.1) and significantly reduced CD86 levels (0.2 ± 0.1) (**Fig. 4I**). In particular, P-60HA/GP accelerated M2 polarization (2.1 ± 0.7), even at early time point.

### P-60HA/GP accelerates bone formation in critical-sized cranial defects

We evaluated the *in vivo* osteogenic capabilities of P-HA scaffolds using a mouse calvarial defect model (**Fig. 5A**). The scaffolds were printed to match the bone defects (4mm×0.3mm), and the new bone formation was monitored by micro-computed tomography (μCT) scanning until 12 weeks (**Fig. 5B**). Considering the similarity in HA content between P-60HA and the natural bone surrounding and the resulting tissue density (**Fig. S7 and S8**), we segmented the area into two regions to facilitate visualization and quantification of bone formation (n=5 for scaffold groups and n=3 for blank). According to the threshold ranges, tissue formations, including the P-60HA scaffold, were highlighted in green and yellow on the μCT images (**Fig. 5B**) and categorized as low-density and high-density immature bone, respectively (**Fig. 5, C and D**). In the lower threshold range (140-300mg HA/cm^3^), the scaffold and soft tissue (brain, scalp, and fat) were disregarded from visualization (**Fig. S8**), allowing focus solely on the tissues infiltrated at the peripheral border and voids of the scaffold. The bone reconstruction process involves infiltration of an ECM network, including the formation of collagen fibrils, followed by mineralization by osteoblasts to create mechanically stable bone in the form of lamellae (*54*). Therefore, the lower threshold range involves the low-density immature bone, including the collagen network and ECM (*54*). The progression of tissue growth and maturation into the high-density threshold range is supported by μCT scanning results taken over time (**Fig. S9**). On the other hand, the higher global threshold range (above 300mg HA/cm^3^) was utilized to evaluate the high-density immature bone, which includes the P-60HA scaffold, unmineralized osteoid, and mineralized tissues.

**Fig. 5.**
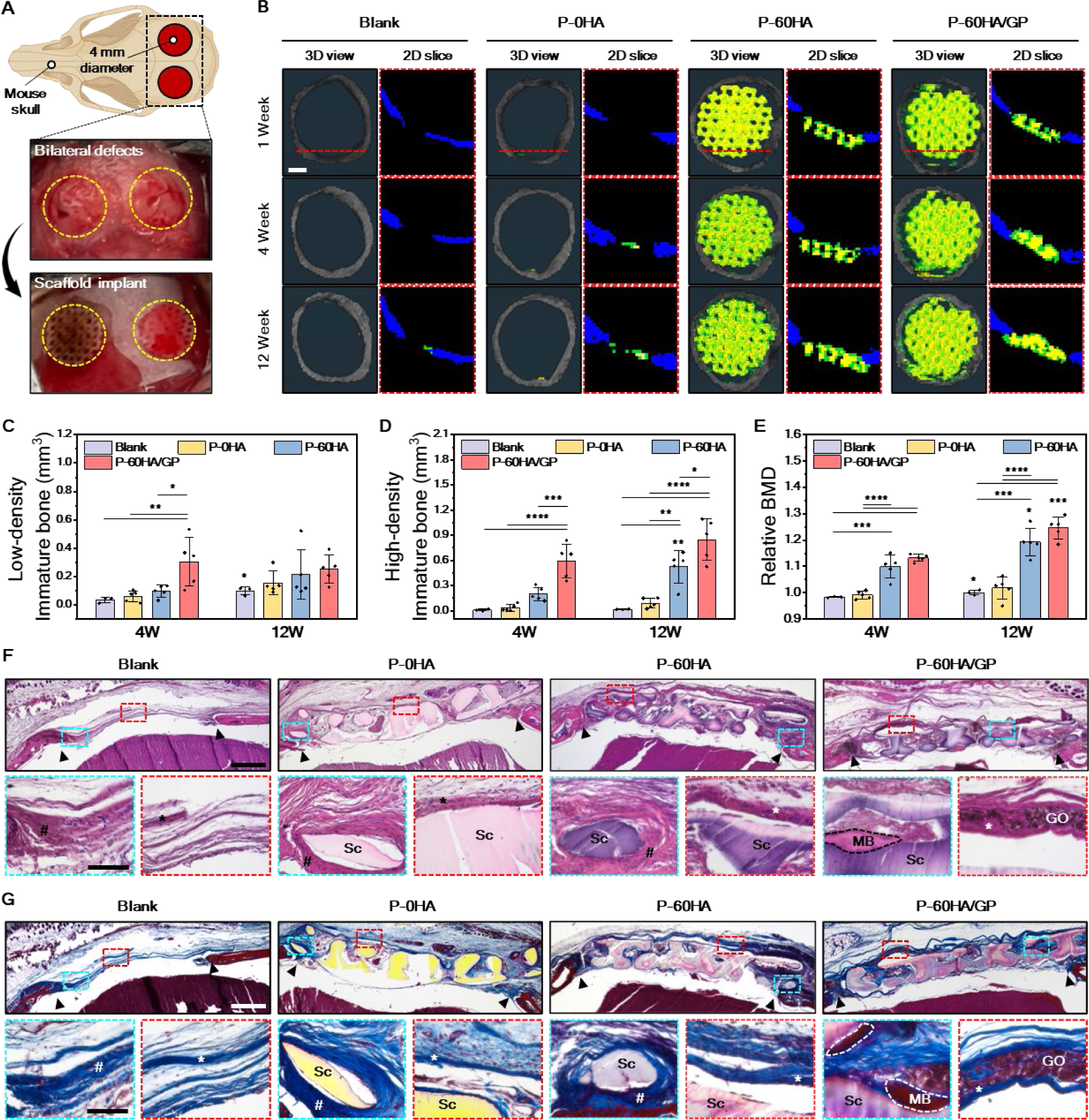
Bone reconstruction by scaffolds in murine critical-sized cranial defect model. (A) Schematic illustration and photos of in vivo cranial defect repair experiment. (B) Representative micro-CT images at 1, 4, and 12 weeks. Blank: no treatment; Green: tissues and scaffolds in the low threshold range; Yellow: tissues and scaffolds in the high threshold range. Scale bar, 1mm. (C and D) Quantitative analyses of immature bone formation in each group at 4 and 12 weeks compared to at 1 week, and (E) relative BMD of each group. \**p* <0.05, \*\**p* <0.01, \*\*\**p* <0.001 and \*\*\*\**p* <0.0001; Error bars, ± SD; n=5. (F) H&E staining and (G) Masson’s trichrome staining of each group at 12 weeks after implantation. Scale bars, 500μm. The images at the bottom represent the higher magnification of each area. Scale bars, 100μm. Sc: scaffold; Black arrow: defective area; Hash (#): unmineralized osteoid; The asterisk (*): periosteal layer; MB: mature bone fragment; GO: GO residue.

The volume of newly formed tissue in the defect area was quantified based on week 1 of each group. P-60HA/GP showed a considerable low-density immature bone formation (0.3 ± 0.2 mm^3^) at an early stage (week 4), exhibiting an 8.4-fold increase compared to blank (0.04 ± 0.02 mm^3^) (\*\**p*<0.01) (**Fig. 5C**). This observation suggests that during the early stage, the presence of GP hydrogel facilitated the development of dense collagen fibers surrounding the scaffold and the formation of ECM found in the outer layer of the scaffold (**Fig. S10**). At 12 weeks, certain experimental groups of P-60HA/GP exhibited a relatively decreased formation compared to week 4. This result indicates maturation in tissue formation due to collagen fiber crosslinking (*54*), transitioning towards the stage of high-density immature bone (**Fig. S10**). Furthermore, within 4 weeks, P-60HA/GP significantly promoted mature bone formation (0.6 ± 0.2 mm^3^), exceeding the blank (0.009 ± 0.01 mm^3^) (\*\*\*\**p* <0.0001) by 69 times and P-60HA (0.2 ± 0.1 mm^3^) (\*\*\**p* <0.001) by 3 times (**Fig. 5D**). In addition, P-60HA/GP enhanced the bone mineral density by 1.2 times at week 12 compared to week 1 (**Fig. 5E and Fig. S7**).

At 12 weeks, the tissues from the center of the defect area were sectioned and examined for analysis. H&E and Masson’s trichrome staining (**Fig. 5, F and G**) showed that P-60HA/GP induced mature new bone fragments between pores and around the structure, while other scaffolds resulted in unmineralized osteoid at the boundary of the defect area. Compared to week 4 (**Fig. S10**), minimal GO residual was observed in the P-60HA/GP implanted group, suggesting gradual *in vivo* degradation of GO (*55*). Moreover, at week 4, P-60HA/GP allowed an abundant tissue network formation throughout its structure, a finding supported by μCT quantification (**Fig. 5C**).

Effective reconstruction of the periosteal layer is crucial to promote bone formation, given its ability to supply abundant growth factors for bone cell growth and differentiation (*56–58*). P- 60HA and P-60HA/GP induced rich periosteal layers around their structure, while the blank and P-0HA formed less connective collagen structures (**Fig. 5G**). Interestingly, collagen fibers were formed along injected GP hydrogel in the P-60HA/GP scaffold, which aligned with the findings from the subcutaneous implantation model (**Fig. 4C**). This outcome demonstrated that P-60HA/GP facilitated the development of aligned collagen formation, which favored bone tissue growth (*56*).

### P-60HA/GP promotes the osteogenesis of endogenous cells and angiogenesis

At week 12, the osteogenic and angiogenic capacity were assessed through IF staining with osteogenic markers (RUNX2, OPN, and OCN) and CD31/α-SMA (**Fig. 6, A and B**). The osteogenic markers were labeled with a red fluorescent dye, and their intensity (n=10 from five biologically independent mice) was evaluated relative to the mean gray value of the blank (n=6 from three biologically independent mice) at week 12 (**Fig. 6C**).

**Fig. 6.**
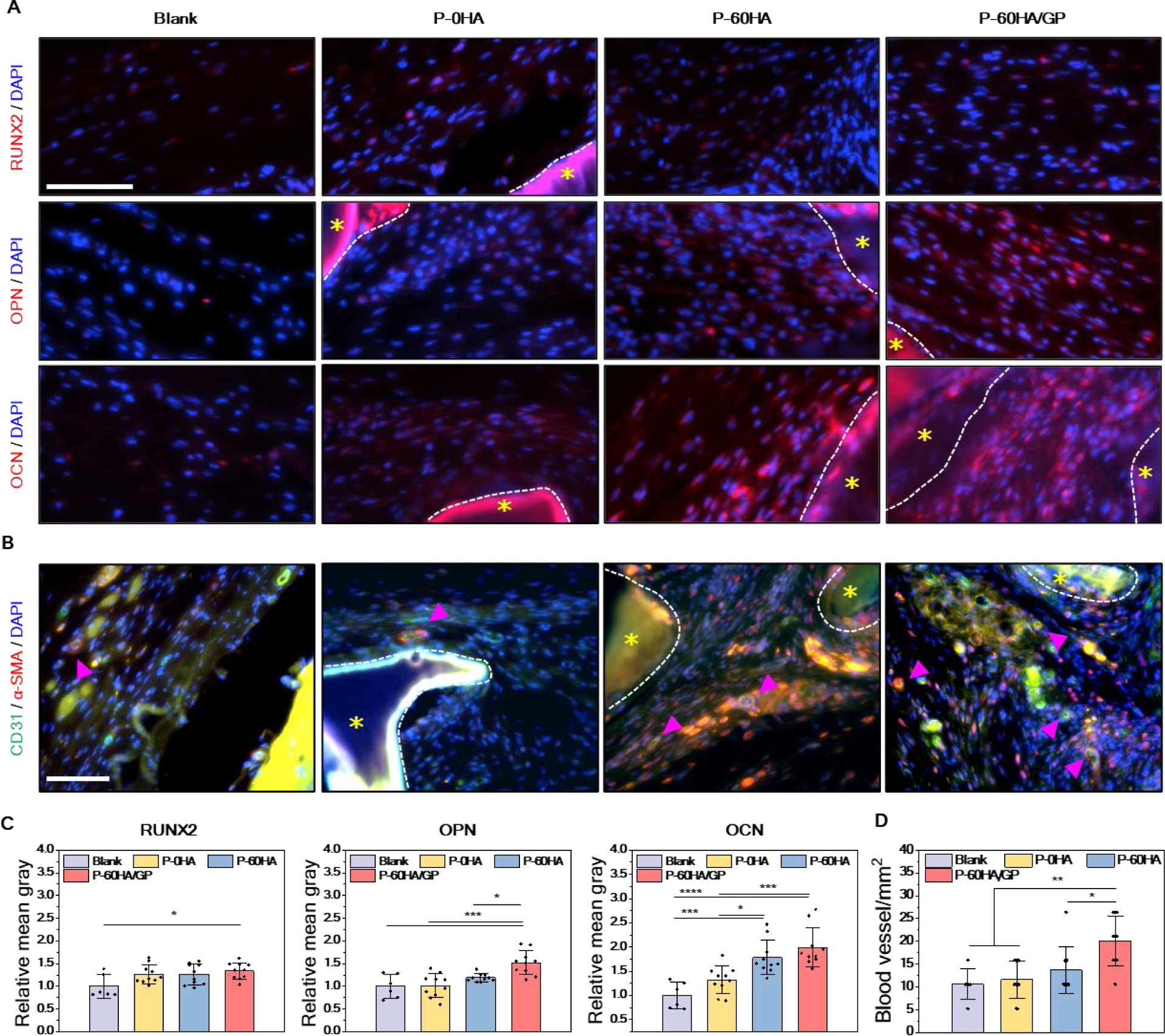
Osteogenic potential and angiogenesis assessments of scaffolds in cranial defect model. (A) Representative immunofluorescence staining images of osteogenic markers and (B) CD31, and α-SMA markers on each scaffold after 12 weeks of implantation. The asterisk (*): scaffold; Pink arrow: newly formed blood vessel. Scale bar, 100μm. (C) Relative mean gray value of each osteogenic differentiation markers on different groups, and (D) the quantitative analysis of blood vessel formation. \**p* <0.05, \*\**p* <0.01, \*\*\**p* <0.001 and \*\*\*\**p* <0.0001; Error bars, ± SD; n=10.

The quantitative results of RUNX2 indicated high expression levels and significant improvements in P-60HA/GP from week 4 (1.5 ± 0.2) (\*\*\**p* <0.001) (**Fig. S11**) to week 12 (1.3 ± 0.2) (\**p* <0.05) compared to the blank, while the other groups exhibited no significant differences (*p* >0.05) from the blank at week 12 (**Fig. 6C**). Furthermore, P-60HA/GP showed strong signal intensities in both OPN (1.5 ± 0.3) (\*\*\**p* <0.001) and OCN (2 ± 0.4) (\*\*\*\**p* <0.0001) compared to those of the blank, which were observed from the early stage (**Fig. S11**). These findings suggest that P-60HA/GP promotes pre-osteoblast proliferation and maturation, thereby facilitating mineralization (*41*).

Significant outcomes were also found in angiogenesis (**Fig. 6, B and D**). While the P- 60HA resulted in slightly higher blood vessel formation (14 ± 5 vessels/mm^2^) compared to the blank (11 ± 3 vessels/mm^2^) and P-0HA (12 ± 4 vessels/mm^2^) groups, this difference was not statistically significant (*p* >0.05). In contrast, P-60HA/GP exhibited a progressive vessel formation (20 ± 5 vessels/mm^2^) (\*\**p* <0.01 for the blank and P-0HA; \**p* <0.05 for P-60HA), particularly pronounced starting from week 4 (12 ± 4 vessels/mm^2^) (**Fig. S12**).

## Discussion

Meeting the global demand for simple and scalable regenerative biomaterials while complying with clinical standards and minimizing cost remains a significant challenge (*59*). The 3D-printed porous CSS presented here provides a rigid (P-60HA)-soft (GP hydrogel) hybrid microenvironment that mimics natural bone (*10*). The fabrication of this hybrid CSS is simple and scalable for manufacturing while meeting the conformal requirements to reconstruct cranial facial defects (*20, 60, 61*). Citrate plays a crucial role as a bioactive factor in bone (*62*). Both mPOC and PPCN, integral components of the CSS, are polymers that belong to a biomaterial technology referred to as citrate-based biomaterials (CBB) (*26, 63*). A CBB, referred to as CITREGEN, has been used for the fabrication of biodegradable implantable medical devices that have been cleared by the U.S. Food and Drug Administration (FDA) to attach soft tissue to bone (*63*). Therefore, given that the FDA is familiar with this new composition and class of polymers (biodegradable thermosets), the CSS is primed for translational application.

Importantly, our strategy eliminates dependence on exogenous biological factors and demonstrates proficient tissue integration and osteogenic potential solely through material-driven cues. The porous architecture of the scaffold contributes to vascularization and tissue ingrowth (*31, 32, 64, 65*). The P-60HA/GP has a heterogeneous pore architecture (**Fig. 2G**) resulting from the printed porous scaffold integrated with the GP hydrogel network. These structures, with their varied pore sizes and interconnected microenvironments, potentially facilitate angiogenesis (*65*) (**Fig. 4F and Fig. 6D**). M2 macrophages play a crucial role in alleviating inflammation and regulating angiogenesis and tissue repair (*53*). Although reactive oxygen species (ROS) are recognized to interfere with M2 activation, they play a role in regulating both pro- and anti-inflammatory macrophage phenotype, depending on the context (*66–68*). PPCN diminishes ROS levels owing to its inherent antioxidant property, reducing oxidative stress (*23*). Meanwhile, GO demonstrates angiogenic activity despite its potential to increase ROS levels dose-dependently (*69*). Moreover, the surface topography resulting from HA particles can influence macrophage polarization (*70*). Hence, P-60HA/GP is expected to have promoted M2 polarization and angiogenesis (**Fig. 4, F and I**) through complex collective actions between components. However, further experiments are warranted to understand the mechanism and their effects.

The osteogenic potential of a biomaterial can be influenced by various factors (*39, 71, 72*). HA encourages osteoblast proliferation and facilitates mesenchymal stem cell growth and differentiation by elevating local Ca^2+^ concentrations (*73, 74*). GO expedites the transformation of stem cells or pre-osteoblasts into osteoblasts by facilitating non-covalent interactions with physiological ions and biomolecules (*43, 75, 76*). Additionally, the role of GO in mineralization, synergizing with HA (*77, 78*), enhances bone formation (*79, 80*). This is supported by a previous report demonstrating that the integration of HA and GO has enhanced osteogenic differentiation in contrast to their individual uses (*79*). The viscoelastic environment of GP hydrogel with rapid stress relaxation promotes intracellular response and tissue remodeling (*37, 39*), and gelatin in GP hydrogel reinforces cell adhesion (*37, 81, 82*). Separately, the carboxylate functional groups of gelatins are expected to have accelerated mineralization through interaction with Ca^2+^ released from HA (*83*). Taken together, these collective properties of P-60HA/GP are believed to promote bone formation and accumulate crucial signaling proteins essential for osteogenesis.

Previous studies demonstrated dose-dependent toxicity of GO through intravenous injection in mice (*84*). A dosage of 0.25mg per mouse from these studies was not toxic or lethal. We administered a 16.5μg GO dose (0.33 mg/mL in 50μL GP hydrogel) within the acceptable tolerance range established by previous studies (*84*). While we demonstrated the potential degradability of GO during *in vivo* bone healing, its long-term effects until complete bone healing are yet unexplored. Further exploration is crucial to determine the optimal GO concentration for osteogenesis and its relationship with biodegradation.

While we have demonstrated the osteogenic capabilities of the CSS, these results have not been directly compared to systems integrating cells and growth factors. However, our primary focus has been on effective bone and tissue reconstruction while minimizing costs, procedures, and potential immune responses, all geared toward swift translation and clinical implementation. Our future work will be expanded to larger animals and will address the potential of our approach for practical applications by comparison in more relevant contexts.

## Ethics approval and consent to participate

The mouse subcutaneous implantation model was carried out with the approval from the Institutional Animal Care and Use Committee at Northwestern University (protocol #IS00003238). The cranial defect animal procedure was performed in compliance with the approval from the University of Chicago Animal Care and Use Committee (ACUP #71745).

## Data and materials availability

All data are available in the main text or the supplementary materials.

## CRediT authorship contribution statement

**Mirae Kim:** Conceptualization, Methodology, Investigation, Visualization, Writing – original draft, Writing – review & editing. **Caralyn P. Collins:** Methodology, Investigation, Visualization, Writing – review & editing. **Yugang Liu**: Methodology, Investigation. **Hsiu-Ming Tsal**: Methodology, Investigation, Visualization. **Yujin Ahn:**Methodology, Investigation, Visualization. **Xinlong Wang:** Methodology, Investigation. **Joseph W. Song**: Methodology, Investigation. **Chongwen Duan:** Methodology, Investigation. **Cheng Sun:** Methodology. **Zhu Yi:** Investigation. **Tong-Chuan He:** Methodology, Writing – review & editing. **Russell R. Reid:** Conceptualization, Investigation, Supervision, Writing – review & editing. **Guillermo A. Ameer:** Conceptualization, Supervision, Writing – review & editing.

## Acknowledgments

This work made use of the IMSERC NMR and Physical Characterization facility at Northwestern University, which has received support from the Soft and Hybrid Nanotechnology Experimental (SHyNE) Resource (NSF ECCS-2025633), and Northwestern University. This work made use of the Keck-II facility and the EPIC facility of Northwestern University’s NUANCE Center, which has received support from the SHyNE Resource (NSF ECCS-2025633), the IIN, and Northwestern’s MRSEC program (NSF DMR-2308691). Imaging work was performed at the Northwestern University Center for Advanced Molecular Imaging (RRID:SCR_021192) generously supported by NCI CCSG P30 CA060553 awarded to the Robert H Lurie Comprehensive Cancer Center.

## References

1. L. A. Kandi, T. L. Jarvis, M. Shrout, D. A. Thornburg, M. A. Howard, M. Ellis, C. M. Teven, Trends in Medicare Reimbursement for the Top 20 Surgical Procedures in Craniofacial Trauma. Journal of Craniofacial Surgery 34, 247–249 (2023).

2. M. J. Carty, N. Ferraro, J. Upton, Reconstruction of pediatric cranial base defects: a review of a single microsurgeon’s 30-year experience. Journal of Craniofacial Surgery 20, 639–645 (2009).

3. S. Bhumiratana, J. C. Bernhard, D. M. Alfi, K. Yeager, R. E. Eton, J. Bova, F. Shah, J. M. Gimble, M. J. Lopez, S. B. Eisig, Tissue-engineered autologous grafts for facial bone reconstruction. Science translational medicine 8, 343ra383-343ra383 (2016).

4. T. W. Bauer, G. F. Muschler, Bone graft materials: an overview of the basic science. Clinical Orthopaedics and Related Research® 371, 10–27 (2000).

5. G. L. Koons, M. Diba, A. G. Mikos, Materials design for bone-tissue engineering. Nature Reviews Materials 5, 584–603 (2020).

6. N. L. Hunter, R. E. Sherman, Combination products: modernizing the regulatory paradigm. Nature Reviews Drug Discovery 16, 513–514 (2017).

7. A. De Pieri, Y. Rochev, D. I. Zeugolis, Scaffold-free cell-based tissue engineering therapies: Advances, shortfalls and forecast. NPJ Regenerative Medicine 6, 18 (2021).

8. F. Shang, Y. Yu, S. Liu, L. Ming, Y. Zhang, Z. Zhou, J. Zhao, Y. Jin, Advancing application of mesenchymal stem cell-based bone tissue regeneration. Bioactive materials 6, 666–683 (2021).

9. J. A. Burdick, R. L. Mauck, J. H. Gorman III, R. C. Gorman, Acellular biomaterials: an evolving alternative to cell-based therapies. Science translational medicine 5, 176ps174- 176ps174 (2013).

10. M. Fontcuberta-Rigo, M. Nakamura, P. Puigbò, Phylobone: a comprehensive database of bone extracellular matrix proteins in human and model organisms. Bone Research 11, 44 (2023).

11. L. Nie, Y. Deng, P. Li, R. Hou, A. Shavandi, S. Yang, Hydroxyethyl chitosan-reinforced polyvinyl alcohol/biphasic calcium phosphate hydrogels for bone regeneration. Acs Omega 5, 10948–10957 (2020).

12. K. Zhou, P. Yu, X. Shi, T. Ling, W. Zeng, A. Chen, W. Yang, Z. Zhou, Hierarchically porous hydroxyapatite hybrid scaffold incorporated with reduced graphene oxide for rapid bone ingrowth and repair. ACS nano 13, 9595–9606 (2019).

13. Z. Zhong, X. Wu, Y. Wang, M. Li, Y. Li, X. Liu, X. Zhang, Z. Lan, J. Wang, Y. Du, Zn/Sr dual ions-collagen co-assembly hydroxyapatite enhances bone regeneration through procedural osteo-immunomodulation and osteogenesis. Bioactive materials 10, 195–206 (2022).

14. P. Song, M. Li, B. Zhang, X. Gui, Y. Han, L. Wang, W. Zhou, L. Guo, Z. Zhang, Z. Li, DLP fabricating of precision GelMA/HAp porous composite scaffold for bone tissue engineering application. Composites Part B: Engineering 244, 110163 (2022).

15. L. Tong, X. Pu, Q. Liu, X. Li, M. Chen, P. Wang, Y. Zou, G. Lu, J. Liang, Y. Fan, Nanostructured 3D-Printed Hybrid Scaffold Accelerates Bone Regeneration by Photointegrating Nanohydroxyapatite. Advanced Science 10, 2300038 (2023).

16. J. Barros, M. P. Ferraz, J. Azeredo, M. Fernandes, P. Gomes, F. Monteiro, Alginate- nanohydroxyapatite hydrogel system: Optimizing the formulation for enhanced bone regeneration. Materials Science and Engineering: C 105, 109985 (2019).

17. M. Kamaraj, A. Datla, S. E. Moulton, S. N. Rath, Biomimetic Mineralization of Mn-Doped Biphasic Calcium Phosphate in the GelMa Hydrogel Acting as a Functional 3D Bioscaffold for Osteo Defect Repair. ACS Applied Polymer Materials 6, 943–955 (2024).

18. G. Kaur, V. Kumar, F. Baino, J. C. Mauro, G. Pickrell, I. Evans, O. Bretcanu, Mechanical properties of bioactive glasses, ceramics, glass-ceramics and composites: State-of-the-art review and future challenges. Materials science and engineering: C 104, 109895 (2019).

19. A. Atala, F. K. Kasper, A. G. Mikos, Engineering complex tissues. Science translational medicine 4, 160rv112-160rv112 (2012).

20. J. R. Tumbleston, D. Shirvanyants, N. Ermoshkin, R. Janusziewicz, A. R. Johnson, D. Kelly, K. Chen, R. Pinschmidt, J. P. Rolland, A. Ermoshkin, Continuous liquid interface production of 3D objects. Science 347, 1349–1352 (2015).

21. R. L. Keate, J. Tropp, C. P. Collins, H. O. T. Ware, A. J. Petty, G. A. Ameer, C. Sun, J. Rivnay, 3D-Printed Electroactive Hydrogel Architectures with Sub-100 µm Resolution Promote Myoblast Viability. Macromolecular bioscience 22, 2200103 (2022).

22. C. G. Jeong, S. J. Hollister, A comparison of the influence of material on in vitro cartilage tissue engineering with PCL, PGS, and POC 3D scaffold architecture seeded with chondrocytes. Biomaterials 31, 4304-4312 (2010).

23. J. Yang, R. Van Lith, K. Baler, R. A. Hoshi, G. A. Ameer, A thermoresponsive biodegradable polymer with intrinsic antioxidant properties. Biomacromolecules 15, 3942–3952 (2014).

24. J. Ye, J. Wang, Y. Zhu, Q. Wei, X. Wang, J. Yang, S. Tang, H. Liu, J. Fan, F. Zhang, A thermoresponsive polydiolcitrate-gelatin scaffold and delivery system mediates effective bone formation from BMP9-transduced mesenchymal stem cells. Biomedical Materials 11, 025021 (2016).

25. M. Liu, X. Zeng, C. Ma, H. Yi, Z. Ali, X. Mou, S. Li, Y. Deng, N. He, Injectable hydrogels for cartilage and bone tissue engineering. Bone research 5, 1–20 (2017).

26. J. Yang, A. R. Webb, G. A. Ameer, Novel citric acid-based biodegradable elastomers for tissue engineering. Advanced Materials 16, 511–516 (2004).

27. M. M. Hasani-Sadrabadi, P. Sarrion, S. Pouraghaei, Y. Chau, S. Ansari, S. Li, T. Aghaloo, A. Moshaverinia, An engineered cell-laden adhesive hydrogel promotes craniofacial bone tissue regeneration in rats. Science translational medicine 12, eaay6853 (2020).

28. H. A. Pearce, J. W. Swain, L. H. Victor, K. J. Hogan, E. Y. Jiang, M. L. Bedell, A. M. Navara, A. Farsheed, Y. S. Kim, J. L. Guo, Thermogelling hydrogel charge and lower critical solution temperature influence cellular infiltration and tissue integration in an ex vivo cartilage explant model. Journal of Biomedical Materials Research Part A 111, 15–34 (2023).

29. A. L. Olivares, È. Marsal, J. A. Planell, D. Lacroix, Finite element study of scaffold architecture design and culture conditions for tissue engineering. Biomaterials 30, 6142–6149 (2009).

30. M. Castilho, A. van Mil, M. Maher, C. H. Metz, G. Hochleitner, J. Groll, P. A. Doevendans, K. Ito, J. P. Sluijter, J. Malda, Melt electrowriting allows tailored microstructural and mechanical design of scaffolds to advance functional human myocardial tissue formation. Advanced Functional Materials 28, 1803151 (2018).

31. W. B. Swanson, M. Omi, Z. Zhang, H. K. Nam, Y. Jung, G. Wang, P. X. Ma, N. E. Hatch, Y. Mishina, Macropore design of tissue engineering scaffolds regulates mesenchymal stem cell differentiation fate. Biomaterials 272, 120769 (2021).

32. V. Karageorgiou, D. Kaplan, Porosity of 3D biomaterial scaffolds and osteogenesis. Biomaterials 26, 5474–5491 (2005).

33. D. D. Deligianni, N. D. Katsala, P. G. Koutsoukos, Y. F. Missirlis, Effect of surface roughness of hydroxyapatite on human bone marrow cell adhesion, proliferation, differentiation and detachment strength. Biomaterials 22, 87–96 (2000).

34. A. B. Faia-Torres, S. Guimond-Lischer, M. Rottmar, M. Charnley, T. Goren, K. Maniura- Weber, N. D. Spencer, R. L. Reis, M. Textor, N. M. Neves, Differential regulation of osteogenic differentiation of stem cells on surface roughness gradients. Biomaterials 35, 9023–9032 (2014).

35. J. W. Song, H. Ryu, W. Bai, Z. Xie, A. Vázquez-Guardado, K. Nandoliya, R. Avila, G. Lee, Z. Song, J. Kim, Bioresorbable, wireless, and battery-free system for electrotherapy and impedance sensing at wound sites. Science Advances 9, eade4687 (2023).

36. C. Zhao, Z. Zeng, N. T. Qazvini, X. Yu, R. Zhang, S. Yan, Y. Shu, Y. Zhu, C. Duan, E. Bishop, Thermoresponsive citrate-based graphene oxide scaffold enhances bone regeneration from BMP9-stimulated adipose-derived mesenchymal stem cells. ACS biomaterials science & engineering 4, 2943–2955 (2018).

37. O. Chaudhuri, L. Gu, D. Klumpers, M. Darnell, S. A. Bencherif, J. C. Weaver, N. Huebsch, H.-p. Lee, E. Lippens, G. N. Duda, Hydrogels with tunable stress relaxation regulate stem cell fate and activity. Nature materials 15, 326–334 (2016).

38. W. R. Legant, J. S. Miller, B. L. Blakely, D. M. Cohen, G. M. Genin, C. S. Chen, Measurement of mechanical tractions exerted by cells in three-dimensional matrices. Nature methods 7, 969–971 (2010).

39. A. K. Gaharwar, I. Singh, A. Khademhosseini, Engineered biomaterials for in situ tissue regeneration. Nature Reviews Materials 5, 686–705 (2020).

40. F. Wang, X. Cai, Y. Shen, L. Meng, Cell–scaffold interactions in tissue engineering for oral and craniofacial reconstruction. Bioactive Materials 23, 16–44 (2023).

41. K. K. Moncal, H. Gudapati, K. P. Godzik, D. N. Heo, Y. Kang, E. Rizk, D. J. Ravnic, H. Wee, D. F. Pepley, V. Ozbolat, Intra-operative bioprinting of hard, soft, and hard/soft composite tissues for craniomaxillofacial reconstruction. Advanced functional materials 31, 2010858 (2021).

42. G. Turnbull, J. Clarke, F. Picard, P. Riches, L. Jia, F. Han, B. Li, W. Shu, 3D bioactive composite scaffolds for bone tissue engineering. Bioactive materials 3, 278–314 (2018).

43. W. C. Lee, C. H. Y. Lim, H. Shi, L. A. Tang, Y. Wang, C. T. Lim, K. P. Loh, Origin of enhanced stem cell growth and differentiation on graphene and graphene oxide. ACS nano 5, 7334–7341 (2011).

44. R. J. Miron, D. D. Bosshardt, Multinucleated giant cells: good guys or bad guys? Tissue Engineering Part B: Reviews 24, 53–65 (2018).

45. C. Yue, C. Ding, X. Du, Y. Wang, J. Su, B. Cheng, Self-assembly of collagen fibrils on graphene oxide and their hybrid nanocomposite films. International Journal of Biological Macromolecules 193, 173–182 (2021).

46. C.-Y. Chen, P.-H. Tsai, Y.-H. Lin, C.-Y. Huang, J. H. Chung, G.-Y. Chen, Controllable graphene oxide-based biocompatible hybrid interface as an anti-fibrotic coating for metallic implants. Materials Today Bio 15, 100326 (2022).

47. H. Bai, C. Li, X. Wang, G. Shi, On the gelation of graphene oxide. The Journal of Physical Chemistry C 115, 5545–5551 (2011).

48. S. Yin, W. Zhang, Z. Zhang, X. Jiang, Recent advances in scaffold design and material for vascularized tissue-engineered bone regeneration. Advanced healthcare materials 8, 1801433 (2019).

49. J. Park, Y. S. Kim, S. Ryu, W. S. Kang, S. Park, J. Han, H. C. Jeong, B. H. Hong, Y. Ahn, B. S. Kim, Graphene potentiates the myocardial repair efficacy of mesenchymal stem cells by stimulating the expression of angiogenic growth factors and gap junction protein. Advanced Functional Materials 25, 2590–2600 (2015).

50. M. H. Norahan, M. Amroon, R. Ghahremanzadeh, M. Mahmoodi, N. Baheiraei, Electroactive graphene oxide-incorporated collagen assisting vascularization for cardiac tissue engineering. Journal of biomedical materials research Part A 107, 204–219 (2019).

51. J. M. Anderson, A. Rodriguez, D. T. Chang, in Seminars in immunology. (Elsevier, 2008), vol. 20, pp. 86-100.

52. R. Sridharan, A. R. Cameron, D. J. Kelly, C. J. Kearney, F. J. O’Brien, Biomaterial based modulation of macrophage polarization: a review and suggested design principles. Materials Today 18, 313–325 (2015).

53. P. Graney, S. Ben-Shaul, S. Landau, A. Bajpai, B. Singh, J. Eager, A. Cohen, S. Levenberg, K. Spiller, Macrophages of diverse phenotypes drive vascularization of engineered tissues. Science advances 6, eaay6391 (2020).

54. W. Wang, K. W. Yeung, Bone grafts and biomaterials substitutes for bone defect repair: A review. Bioactive materials 2, 224–247 (2017).

55. C. M. Girish, A. Sasidharan, G. S. Gowd, S. Nair, M. Koyakutty, Confocal Raman imaging study showing macrophage mediated biodegradation of graphene in vivo. Advanced healthcare materials 2, 1489–1500 (2013).

56. J. Pan, H. Li, K. Jin, H. Jiang, K. Li, Y. Tang, Z. Liu, K. Zhang, K. Chen, Z. Xu, Periosteal topology creates an osteo-friendly microenvironment for progenitor cells. Materials Today Bio 18, 100519 (2023).

57. S. J. Roberts, N. Van Gastel, G. Carmeliet, F. P. Luyten, Uncovering the periosteum for skeletal regeneration: the stem cell that lies beneath. Bone 70, 10–18 (2015).

58. R. Tevlin, M. Griffin, K. Chen, M. Januszyk, N. Guardino, A. Spielman, S. Walters, G. E. Gold, C. K. Chan, G. C. Gurtner, Denervation during mandibular distraction osteogenesis results in impaired bone formation. Scientific Reports 13, 2097 (2023).

59. A. E. Jakus, A. L. Rutz, S. W. Jordan, A. Kannan, S. M. Mitchell, C. Yun, K. D. Koube, S. C. Yoo, H. E. Whiteley, C.-P. Richter, Hyperelastic “bone”: A highly versatile, growth factor–free, osteoregenerative, scalable, and surgically friendly biomaterial. Science translational medicine 8, 358ra127-358ra127 (2016).

60. M. Zhang, R. Lin, X. Wang, J. Xue, C. Deng, C. Feng, H. Zhuang, J. Ma, C. Qin, L. Wan, 3D printing of Haversian bone–mimicking scaffolds for multicellular delivery in bone regeneration. Science advances 6, eaaz6725 (2020).

61. X. Xue, Y. Hu, S. Wang, X. Chen, Y. Jiang, J. Su, Fabrication of physical and chemical crosslinked hydrogels for bone tissue engineering. Bioactive materials 12, 327–339 (2022).

62. J. Guo, X. Tian, D. Xie, K. Rahn, E. Gerhard, M. L. Kuzma, D. Zhou, C. Dong, X. Bai, Z. Lu, Citrate-based tannin-bridged bone composites for lumbar fusion. Advanced Functional Materials 30, 2002438 (2020).

63. H. Wang, S. Huddleston, J. Yang, G. A. Ameer, Enabling Proregenerative Medical Devices via Citrate-Based Biomaterials: Transitioning from Inert to Regenerative Biomaterials. Advanced Materials, 2306326 (2023).

64. X. Xiao, W. Wang, D. Liu, H. Zhang, P. Gao, L. Geng, Y. Yuan, J. Lu, Z. Wang, The promotion of angiogenesis induced by three-dimensional porous beta-tricalcium phosphate scaffold with different interconnection sizes via activation of PI3K/Akt pathways. Scientific reports 5, 9409 (2015).

65. H. Mehdizadeh, S. Sumo, E. S. Bayrak, E. M. Brey, A. Cinar, Three-dimensional modeling of angiogenesis in porous biomaterial scaffolds. Biomaterials 34, 2875–2887 (2013).

66. S. Gordon, P. R. Taylor, Monocyte and macrophage heterogeneity. Nature reviews immunology 5, 953–964 (2005).

67. H.-Y. Tan, N. Wang, S. Li, M. Hong, X. Wang, Y. Feng, The reactive oxygen species in macrophage polarization: reflecting its dual role in progression and treatment of human diseases. Oxidative medicine and cellular longevity 2016, (2016).

68. Y. Zhang, S. Choksi, K. Chen, Y. Pobezinskaya, I. Linnoila, Z.-G. Liu, ROS play a critical role in the differentiation of alternatively activated macrophages and the occurrence of tumor-associated macrophages. Cell research 23, 898–914 (2013).

69. S. Mukherjee, P. Sriram, A. K. Barui, S. K. Nethi, V. Veeriah, S. Chatterjee, K. I. Suresh, C. R. Patra, Graphene oxides show angiogenic properties. Advanced healthcare materials 4, 1722–1732 (2015).

70. Y. Zhu, H. Liang, X. Liu, J. Wu, C. Yang, T. M. Wong, K. Y. Kwan, K. M. Cheung, S. Wu, K. W. Yeung, Regulation of macrophage polarization through surface topography design to facilitate implant-to-bone osteointegration. Science advances 7, eabf6654 (2021).

71. A. J. Engler, S. Sen, H. L. Sweeney, D. E. Discher, Matrix elasticity directs stem cell lineage specification. Cell 126, 677–689 (2006).

72. B. Conrad, C. Hayashi, F. Yang, Gelatin-based microribbon hydrogels support robust MSC osteogenesis across a broad range of stiffness. ACS biomaterials science & engineering 6, 3454–3463 (2020).

73. H. Shi, Z. Zhou, W. Li, Y. Fan, Z. Li, J. Wei, Hydroxyapatite based materials for bone tissue engineering: A brief and comprehensive introduction. Crystals 11, 149 (2021).

74. D. G. Castner, B. D. Ratner, Proteins controlled with precision at organic, polymeric, and biopolymer interfaces for tissue engineering and regenerative medicine. Principles of Regenerative Medicine, 523–534 (2019).

75. A. M. Arnold, B. D. Holt, L. Daneshmandi, C. T. Laurencin, S. A. Sydlik, Phosphate graphene as an intrinsically osteoinductive scaffold for stem cell-driven bone regeneration. Proceedings of the National Academy of Sciences 116, 4855–4860 (2019).

76. W. Zhang, Q. Chang, L. Xu, G. Li, G. Yang, X. Ding, X. Wang, D. Cui, X. Jiang, Graphene oxide-copper Nanocomposite-coated porous CaP scaffold for vascularized bone regeneration via activation of Hif-1α. Advanced healthcare materials 5, 1299–1309 (2016).

77. C. Santos, C. Piedade, P. Uggowitzer, M. Montemor, M. Carmezim, Parallel nano- assembling of a multifunctional GO/HapNP coating on ultrahigh-purity magnesium for biodegradable implants. Applied Surface Science 345, 387–393 (2015).

78. Q. Wang, Y. Chu, J. He, W. Shao, Y. Zhou, K. Qi, L. Wang, S. Cui, A graded graphene oxide-hydroxyapatite/silk fibroin biomimetic scaffold for bone tissue engineering. Materials Science and Engineering: C 80, 232–242 (2017).

79. J. H. Lee, Y. C. Shin, S.-M. Lee, O. S. Jin, S. H. Kang, S. W. Hong, C.-M. Jeong, J. B. Huh, D.-W. Han, Enhanced osteogenesis by reduced graphene oxide/hydroxyapatite nanocomposites. Scientific reports 5, 18833 (2015).

80. W. Nie, C. Peng, X. Zhou, L. Chen, W. Wang, Y. Zhang, P. X. Ma, C. He, Three- dimensional porous scaffold by self-assembly of reduced graphene oxide and nano- hydroxyapatite composites for bone tissue engineering. Carbon 116, 325–337 (2017).

81. J. M. Aamodt, D. W. Grainger, Extracellular matrix-based biomaterial scaffolds and the host response. Biomaterials 86, 68–82 (2016).

82. K. H. Vining, D. J. Mooney, Mechanical forces direct stem cell behaviour in development and regeneration. Nature reviews Molecular cell biology 18, 728–742 (2017).

83. X. Wu, T. Zhang, B. Hoff, S. Suvarnapathaki, D. Lantigua, C. McCarthy, B. Wu, G. Camci- Unal, Mineralized hydrogels induce bone regeneration in critical size cranial defects. Advanced Healthcare Materials 10, 2001101 (2021).

84. K. Wang, J. Ruan, H. Song, J. Zhang, Y. Wo, S. Guo, D. Cui, Biocompatibility of graphene oxide. Nanoscale Res Lett 6, 1–8 (2011).

