## Supplementary Material for "Personalized composite scaffolds for accelerated cell- and growth factor-free craniofacial bone regeneration"

### Supplementary Materials

**The file includes:**

Materials and Methods

Figs. S1 to S12

Table S1

### Materials and Methods

#### Synthesis of mPOC polymer and preparation of P-HA composite

mPOC polymer was synthesized by a two-step process: (1) the synthesis of POC pre-polymer and (2) the conjugation of methacrylate groups onto the pre-polymer. First, equal molar amounts of citric acid and 1,8-octanediol were added to a round-bottom flask and melted at 165°C. The mixture was then transferred into a 140°C oil bath and continuously stirred for 55 minutes under nitrogen gas. Subsequently, the mixture was purified and precipitated with MQ water and lyophilized to obtain the pre-polymer. Following lyophilization, the POC pre-polymer (66g) was dissolved in tetrahydrofuran (THF, 540mL) at 60°C, and imidazole was added once the mixture became clear. Glycidyl methacrylate was slowly added and allowed to react for 6 hours under reflux. After completion of the reaction, the mixture was concentrated using a rotary evaporator, and the crude product was obtained by precipitation in excess MQ water. After purifying with MQ water, the final product was obtained by lyophilization. Characterization of the product was performed using FT-IR (Thermo Nicolet Nexus, ART mode) and $^{1}$H-NMR (X500) spectroscopy.

To prepare the P-HA composite, mPOC polymer and HA powder were mixed in various ratios while maintaining an overall concentration of 70wt.% in pure ethanol. The mPOC-to-HA ratios used were 100:0, 80:20, 60:40, and 40:60. After thorough mixing, Irgacure 819 and Ethyl 4-dimethylamino benzoate (EDAB) were added to the mixture at a concentration of 3wt.% as co-photoinitiators.

**Fabrication of the 3D-printed P-HA scaffolds**

A custom-built micro-continuous liquid interface production printer was utilized to fabricate these samples. CAD files were first designed in SolidWorks to create the model for printing. Relevant parameters of this model were a scaffold diameter of 5.125mm, and height of 0.5mm, and thickness of 125µm for *in vitro* experiments and a scaffold diameter of 4mm, height of 0.3mm, and strut thicknesses of 125µm for the cranial defect experiments. These models were subsequently sliced into cross-sectional images via the use of a homemade MATLAB code with 5µm layer thickness. The resulting cross-sectional images could then be projected onto the resin bath via the use of a digital micromirror device (DMD) with a wavelength of 365nm. The lateral pixel resolution during printing was 7.1µm x 7.1µm. Power delivery for each layer of the part was 0.948 mJ/cm2, 1.608 mJ/cm2, 2.092 mJ/cm2, and 2.552 mJ/cm2 for the 0% HA, 20% HA, 40% HA, and 60% HA composite, respectively. Teflon was used as the material for the oxygen permeable membrane; this resulted in an oxygen-rich region at the bottom of the resin bath, which prevented polymerization of the resin along the membrane surface and allowed for a continuous printing process.

**Characterization of P-HA scaffolds**

The mechanical properties of P-HA structures were assessed by a universal testing machine (Model 5940, Instron, High Wycombe, UK). They were printed in a plug structure with 3mm diameter and 6mm height dimensions. The samples were immersed in PBS at RT overnight before the measurement. The compressive crosshead displacement was applied to the plug at 2mm $min^{-1}$

rate until it reached the breaking point, and the stress versus strain curve was obtained. The compressive modulus was determined by calculating the slope of the stress-strain curve within the initial 10% strain range.

The degradation kinetics of P-HA scaffolds were evaluated by monitoring changes in mass over time. The scaffolds were immersed in PBS at 80°C, and mass loss was recorded weekly for 13 weeks. The degradation behavior was normalized to the initial mass and then represented as a mass versus time curve.

The structural morphology was visualized using scanning electron microscopy (SEM) (S-4800, Hitachi, Japan) and sputter-coated with gold and platinum for analysis. The samples were rapidly quenched with liquid nitrogen immediately after being incorporated with the GP hydrogel to preserve the hydrogel structure. Subsequently, the samples were freeze-dried for the analysis.

**Synthesis of PPCN polymer**

PPCN was obtained by following the previous studies. Briefly, Poly(polyethylene glycol citrate) acrylate (PPCac) pre-polymer was synthesized by melting citric acid, polyethylene glycol (Mw 400), and 1,3-diglycerolate diacrylate with the feed molar ratio of 5:9:1, and stirred at 140°C for 45 minutes under nitrogen gas. The obtained PPCac pre-polymer and NIPAAm were added to a single system in a 1:1 molar ratio and dissolved in 1,4-dioxane at 65°C. Azobis-(isobutyronitrile) (AIBN), radical initiator, was added to the mixture and reacted overnight in a nitrogen atmosphere. After the reaction, the mixture was diluted in 1,4-dioxane, and the product was obtained by precipitating in diethyl ether and vacuum-dried. The product was then neutralized to pH 7.4 and dialyzed (2kDa MWCO) against MQ water at 4°C and lyophilized. The chemical structure of PPCN was analyzed with $^1$H-NMR (X500) and FT-IR (Thermo Nicolet Nexus) spectroscopy.

**Fabrication of GO-PPCNg hydrogel composite**

To prepare GP hydrogel composite precursor, PPCN powder was initially solubilized in PBS at a concentration of 100mg/mL. The PPCN stock solution was then diluted at a 1:1 ratio with a gelatin solution (2mg/mL in PBS) to obtain the PPCNg solution. Subsequently, the graphene oxide solution (2mg/mL, Sigma-Aldrich) was introduced into the PPCNg solution at a volume ratio of 1:5, and thorough mixing was performed on ice. The resulting GP precursor solution was then stored at 4°C until it was ready for use.

**Characterization of GO-PPCNg hydrogel composite**

The rheological properties of the thermoresponsive hydrogels were evaluated using a Discovery Hybrid DHR-3 (TA, New Castle, DE). Each hydrogel precursor solution was placed in a parallel plate geometry with a 1mm gap, and a solvent trap filled with water was applied to prevent water loss during the experiment. The changes in storage moduli (G′) and loss moduli (G″) were measured in oscillatory temperature ramp mode (1°C/min) over the range of 15°C to 45°C. The tests were performed at a constant frequency of 1Hz and a strain of 1%. The gelation temperature was determined by the point at which G′ and G″curves cross over.

The stress relaxation was evaluated by using Discovery Hybrid DHR-3 (TA, New Castle, DE). The hydrogel precursor solution was prepared in ice and applied onto the plate of the rheometer at 37°C. The gel was covered with parallel plate geometry with a 1mm gap and allowed to sit for about 1 min until the stress stabilized. Once the initial equilibrium was reached, stress relaxation was performed under a consistent strain of 15% at a frequency of 1Hz.

***In vitro* cell compatibility assessment**

The bone marrow-derived mesenchymal stem cells (hMSC) (PCS-500-012) sourced from the American Type Culture Collection (ATCC) were used for the *in vitro* experiments. They were cultured in a growth medium (PCS-500-030, ATCC) supplemented with growth kit (PCS-500-040, ATCC) and incubated at 37°C in a 5% $CO_2$ environment. Prior to cell seeding, all samples were sterilized by an ethylene oxide sterilization system, and the scaffolds were immersed in the growth media for 24 hours at 37°C.

For the assessment of biocompatibility, hMSCs (passage 5) were seeded at a density of 20,000 cells per well in 48-well plates. Subsequently, the scaffolds (5mmD × 0.5mmH) were placed in each well after 30 minutes of seeding. For the mPOC-HA/GP composite scaffold, GP hydrogel (50μL) precursor solution was applied to the scaffold. The growth medium was refreshed every 48 hours, and alamarBlue assay and Live/dead staining were performed at the desired time point following the manufacturer's instruction.

Confocal microscopy was used to illustrate cytoskeleton staining. 10μL of hMSCs suspension with 40,000 cells were seeded on the scaffold, and the growth medium was added in each well after 30 minutes. After 4 days of culture, the cells-scaffolds were fixed with 4% paraformaldehyde and stained with 4′,6-diamidino-2-phenylindole (DAPI), and rhodamine-phalloidin to visualize cell nuclei and F-actin, respectively.

***In vitro* osteogenic differentiation of hMSCs**

The low passage number of hMSCs (below passage 4) was used to assess the osteogenic expression in the following experiments. The cells were prepared in the growth medium for 48 hours, and the medium was changed to the osteocyte differentiation medium (PCS-500-052, ATCC).

The intracellular alkaline phosphatase (ALP) activity was conducted using the ALP assay kit (K422-500, Biovision), following the manufacturer's instructions. The cell lysate was collected using a lysate buffer solution, followed by the addition of an equal volume of cell lysate to individual wells of a 96-well plate. 5mM p-nitrophenyl phosphate was then introduced each well. After 1 hour of incubation in the dark, the optical density (OD) value was measured at 405nm using a citation-5 imaging reader (BioTek).

To normalize the ALP activity, the Quanti-iT PicoGreen dsDNA assay (Invitrogen) was used to determine the total DNA concentration in each sample, following the provided instructions. The cell lysate was mixed with the working solution and incubated in a 96-well plate, protected from light. The fluorescent signal was detected after 5 minutes with excitation at 480nm and emission at 520nm, using a citation-5 imaging reader.

Real-time reverse quantitative PCR (RT-qPCR) was performed to analyze the relative gene expression level of osteogenic markers at 7, 14, and 21 days of culture. The RNA isolation from the sample was obtained using the Aurum Total RNA mini-Kit (Bio-Rad). The purification of isolated RNA was conducted according to the manual, and the yield was determined using the citation 5 imaging reader. The RT-qPCR process was conducted using iTaq Universal Sybr Green One-step Kit (Bio-Rad) in CFX96 (Bio-Rad) system. The designed primer sequences were described in the Supplementary table (**Table S1**). The gene expression levels were quantified using the $2^{(-\Delta\Delta CT)}$ method and normalized to the value of GAPDH housekeeping gene, followed by additional normalization to the value of P-0HA at day 0 to facilitate comparison.

***In vivo* biocompatibility test through mouse subcutaneous implantation**

C57BL/6 mice (8 to 10 weeks, male) were purchased from the Jackson Laboratory (Bar Harbor, ME, USA). For this study, three mice were allocated to each group at different time points. In the procedure, incisions were made on each side of the back. The scaffolds from each experimental group were implanted in each mouse (three mice at each time pointe), resulting in a total of 6 implants per group (n=6). Implants were harvested with the tissues for the examination of the immune response and cell/tissue infiltration at the time points of day 7 and day 35.

For the histological analysis, the harvested tissues were fixed using 4% paraformaldehyde and immersed in 10% EDTA solution. The decalcifying solution was changed every 2 days under continuous stirring at 4°C for 2 weeks.

***In vivo* cranial defect reconstruction**

C57BL/6 mice (8 weeks, male) were purchased from Harlan Envigo (Indianapolis, IN). Mice were anesthetized with isoflurane during the procedure, and a 4mm diameter of bilateral defect was created on the cranial bone of each mouse. Subsequently, the pericranium was removed by scraping to completely expose the calvaria. The scaffold was implanted on the defect, while the blank group was left without any implant. For the P-60HA/GP composite scaffold, 50μL of GP hydrogel was further introduced. Following implantation, a thin mPOC film was applied on the site with a small amount of thrombin/fibrinogen hydrogel to secure the implant. The progress of bone reconstruction was monitored via micro-CT scanning (X-CUBE) throughout the experimental period. Upon completion of the experiment, after sacrificing the mice, the skull samples were collected at 4 weeks and 12 weeks. These skull samples were subsequently fixed and decalcified with Cal-EX II (Fisher Scientific) for a duration of 48 hours in preparation for the histological analysis.

### Micro-CT scanning and analysis

The imaging was performed using the X-CUBE system (Molecubes NV, Gent, Belgium). The acquisition was carried out using the spiral high-resolution protocol in an X-ray source with settings at 50kVp and 440μA. For the reconstruction of volumetric CT images, an iterative image space reconstruction algorithm was applied. The resulting images were reconstructed in a 400 × 400 × 370 format, with each voxel having a size of 100 × 100 × 100μm³.

An Amira software (Thermo Scientific) was used for the 3D reconstruction and the tissue formation analysis. The P-60HA scaffold exhibited bone mineral density (BMD) similar to native bone, and it was visible across a broad range of thresholds. Therefore, the assessment of tissue formation was conducted in different threshold ranges. In the lower global threshold region (140 to 300mg HA/cm$^3$), where neither soft tissue nor P-60HA scaffolds were visible, the low-density immature bone was quantified. A higher global threshold (above 300mg HA/cm$^3$) was applied to visualize both the P-60HA scaffold and native bone and to quantify the high-density immature bone. The volume observed in week 1 was used as the baseline to calculate the newly formed volume.

BMD was investigated by segmenting the defective area in the micro-CT images of each group using the ImageJ Fiji (NIH) program. BMD was assessed by comparing the ratio of values obtained at week 12 to those at week 1.

### Tissue processing and histological analysis

For histological evaluation, all tissue samples were fixed using 4% paraformaldehyde and decalcified after harvesting. Tissue samples were then dehydrated and embedded in paraffin for the following experiments.

The tissue specimens were sectioned to 10μm thickness and stained with Hematoxylin and eosin (H&E) and Masson's trichrome (Masson Trichome Kit 87019, Epredia) to evaluate the biocompatibility and the tissue formation. The stained tissues were examined under a bright-field microscope (Eclipse Ti2-E, Nikon).

**Immunofluorescence (IF) staining**

For the IF staining, the tissue sections were treated with heat induced epitope retrieval and blocked with a 1% bovine serum albumin solution at room temperature for 30 minutes. Following the blocking step, the sections were incubated overnight at 4°C with primary antibodies, CD86 (1:200, 14-0862-82, Invitrogen), CD163 (1:500, ab182422, Abcam), F4/80 (1:50, SC-25830, Santa Cruz Biotech.), CD31 (1:200, ab182981, Abcam), α-Smooth Muscle Actin (α-SMA) (1:1000, ab7817, Abcam), RUNX2 (1:50, SC-390351, Santa Cruz.), Osteopontin (OPN) (1:50, SC-21742, Santa Cruz Biotech.), and Osteocalcin (OCN) (1:100, 23418-1-AP, Proteintech.). To visualize the immunofluorescence labeling, Alexa Fluor 488 anti-mouse (1:1000, Invitrogen), Fluor 488 anti-rabbit (1:1000, Invitrogen), and Alexa Fluor 594 anti-rabbit (1:1000, Invitrogen) secondary antibodies were used, with DAPI for nuclei. The sections were scanned by microscope (Eclipse Ti2-E, Nikon) with Imaging software (NIS-Elements, Nikon). Relative mean gray from IF staining was analyzed using the ImageJ program.

**Statistical analysis**

All the results were presented as mean value ± standard deviation (SD). The mechanical properties and *in vitro* assessments were plotted based on n=3. In the subcutaneous implantation model, each group was assessed using n=6, sourced from three biologically independent mice. For the cranial defect model, n=5 mice per group (or n=3 for the blank) were utilized for CT quantification, while n=10 (or n=6 for the blank), obtained from five (or three for the blank) biologically independent mice in each group, were employed for IF staining analysis. Statistical analyses were determined by a one-way ANOVA with Tukey's post hoc test to compare three or more groups. Significance values were shown with three levels. $^{*}p < 0.05$, $^{**}p < 0.1$, $^{***}p < 0.001$, and $^{****}p < 0.0001$.

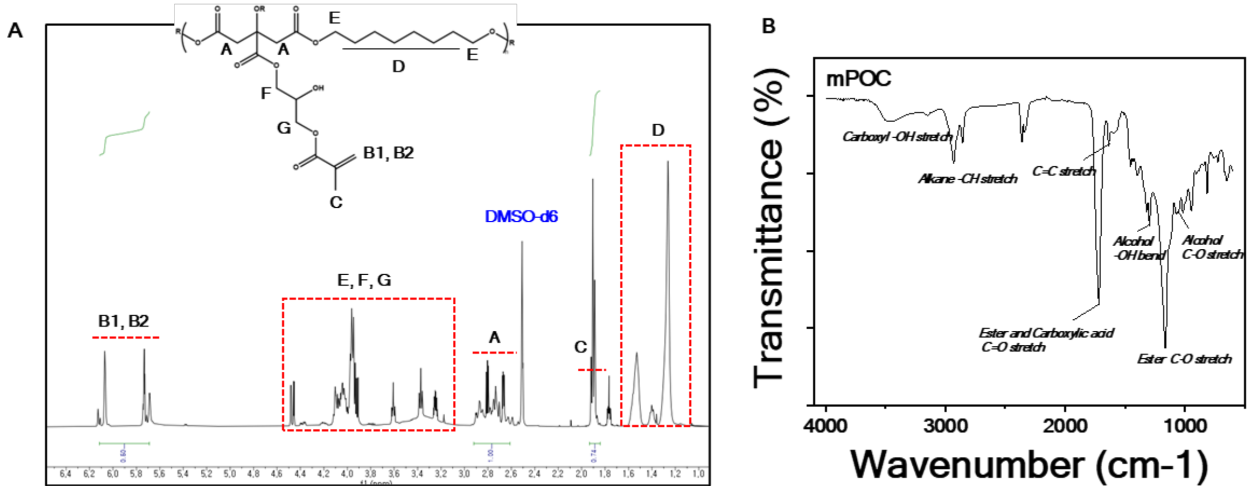


**Fig. S1. Representative $^1$H-NMR spectra and FT-IR spectra of mPOC polymer.** (**A**) Characteristic peaks corresponding to POC, and methacrylate group are assigned in $^1$H-NMR spectra. (**B**) In FT-IR spectra, various functional groups within the mPOC polymer are assigned.

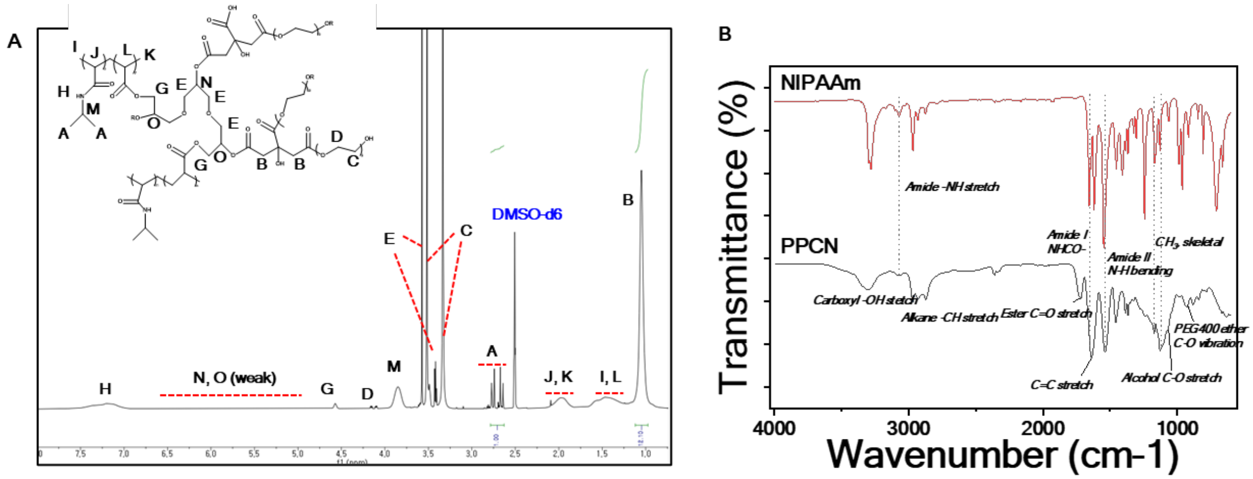


**Fig. S2. Representative $^{1}$H-NMR spectra and FT-IR spectra of PPCN polymer**. (**A**) The $^{1}$H-NMR spectra identify and assign characteristic peaks corresponding to PNIPAAm, PEG, and citric acid are assigned. (**B**) Identification and characterization of different functional groups within PNIPAAm and PPCN polymers by FT-IR analysis.

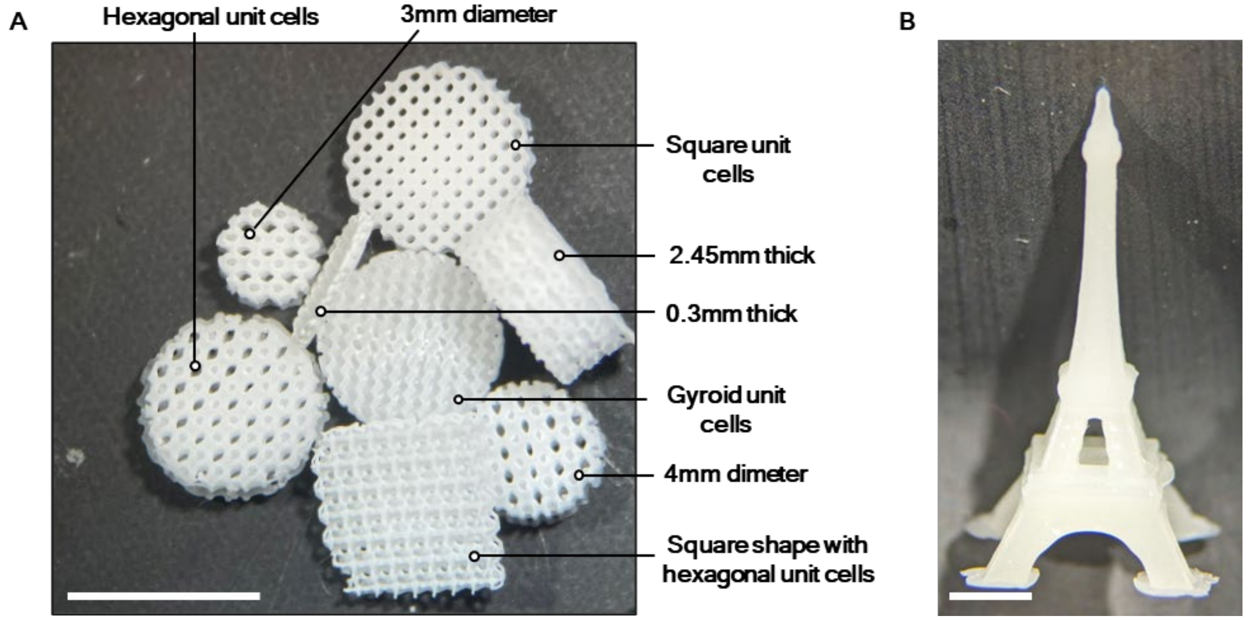


**Fig. S3. 3D-printed P-HA structures with μCLIP 3D printer.** (**A**) 3D-printed porous P-60HA scaffolds with various types of unit cells and dimensions. Scale bar, 5mm. (**B**) 3D-printed Eiffel Tower with P-HA. Scale bar, 2mm.

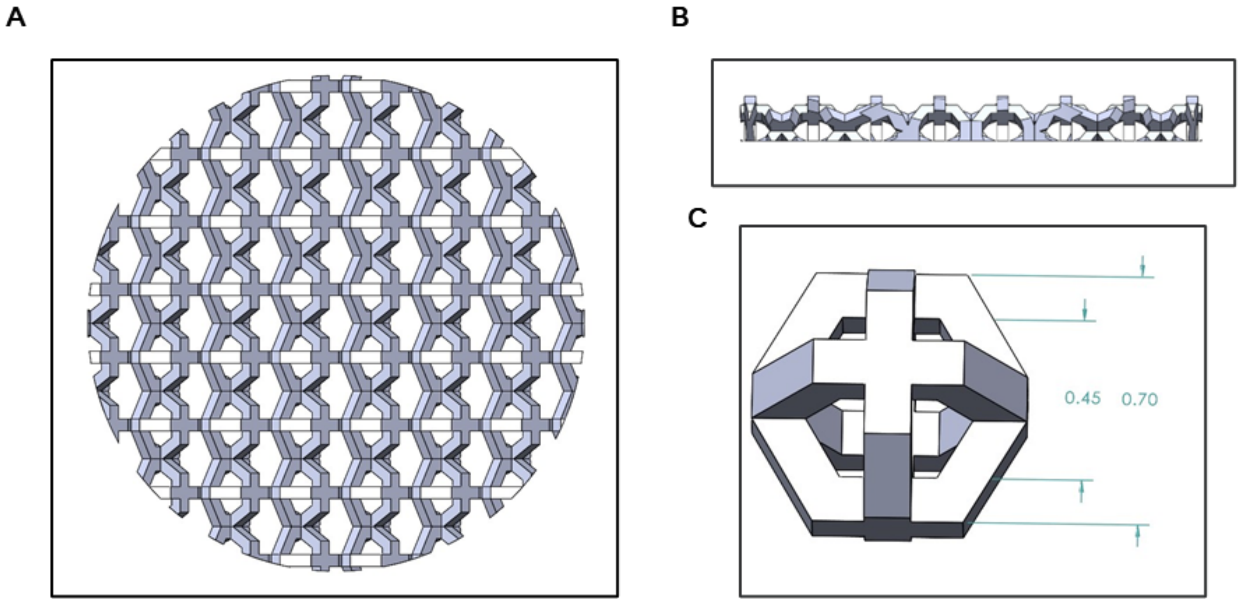


**Fig. S4. 3D design diagram of a porous scaffold with hexagonal unit cells.** (**A**) Top view and (**B**) side view of the designed structure. The diameter and height of the scaffold were adjusted according to the experiment design. (**C**) Representative view of the unit cells in 3D-designed structure and its dimension. Unit: mm.

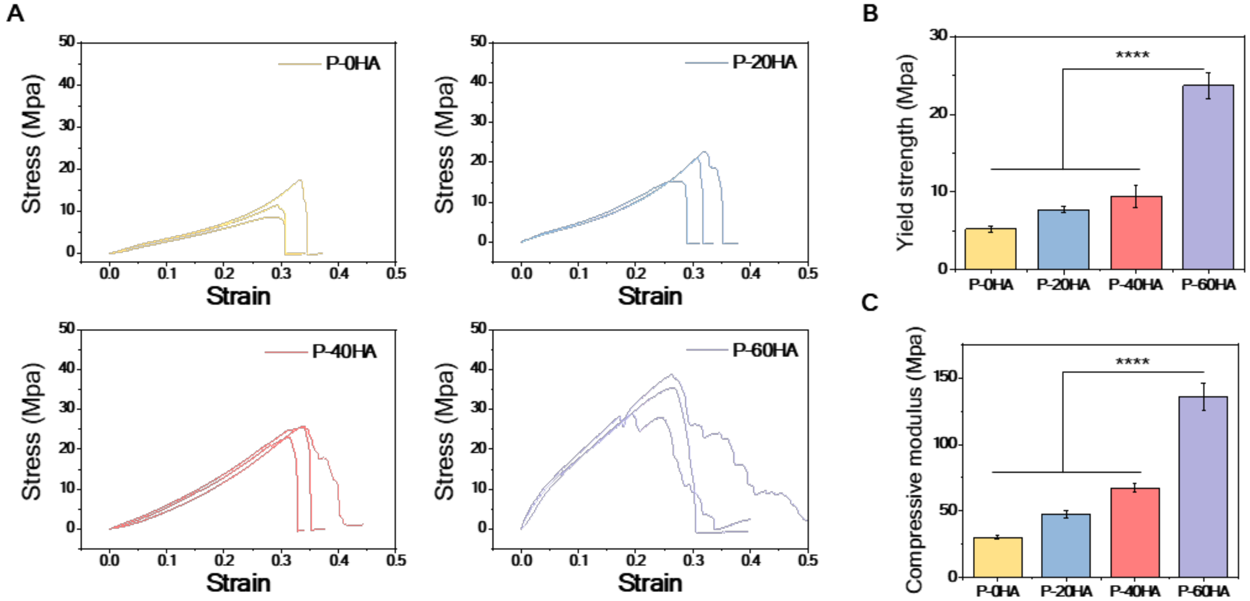


**Fig. S5. The assessment of mechanical properties of P-HA groups under compressive load.** (**A**) Stress-strain curves of P-HA groups. (**B and C**) The calculated compressive strength and compressive moduli of P-HA groups within the initial 10% strain range. $^{****}p < 0.0001$; Error bars, ± SD; n=3.

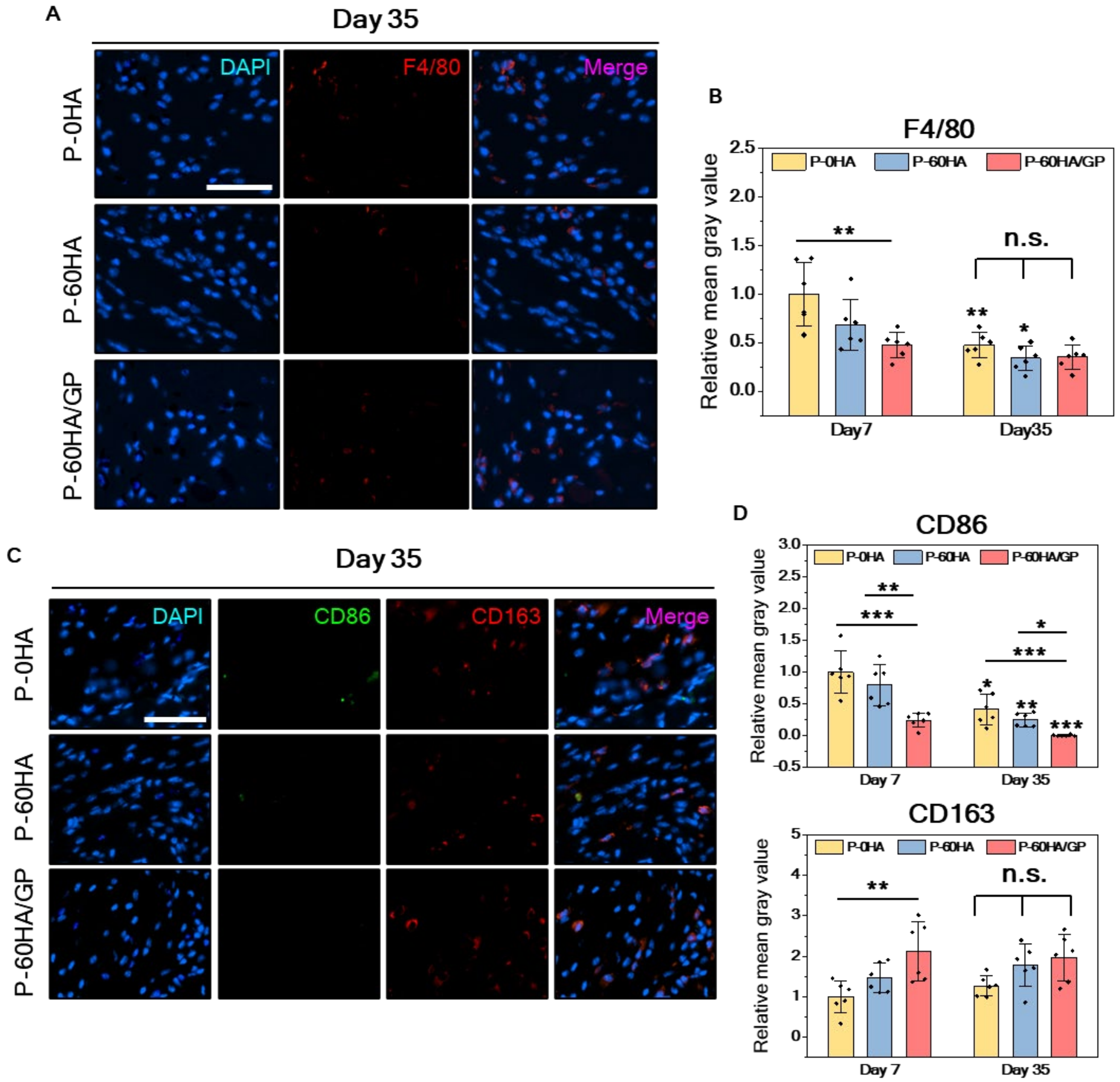


Fig. S6. **Representative immunofluorescence images of F4/80, CD86, and CD163 markers along with quantitative analyses conducted at 35 days post-implantation.** (**A**) F4/80 marker at day 35 for each sample, and (**B**) quantitative analysis of each group in comparison to day 7. (**C**) Representative images of CD86 and CD163 markers at day 35, and (**D**) quantitative result of each marker at different time points. Scale bars, 50μm. $^{*}p$ <0.05, $^{**}p$ <0.01, $^{***}p$ <0.001 and $^{****}p$ <0.0001; n.s.: no significant difference; Error bars, ± SD; n=6.

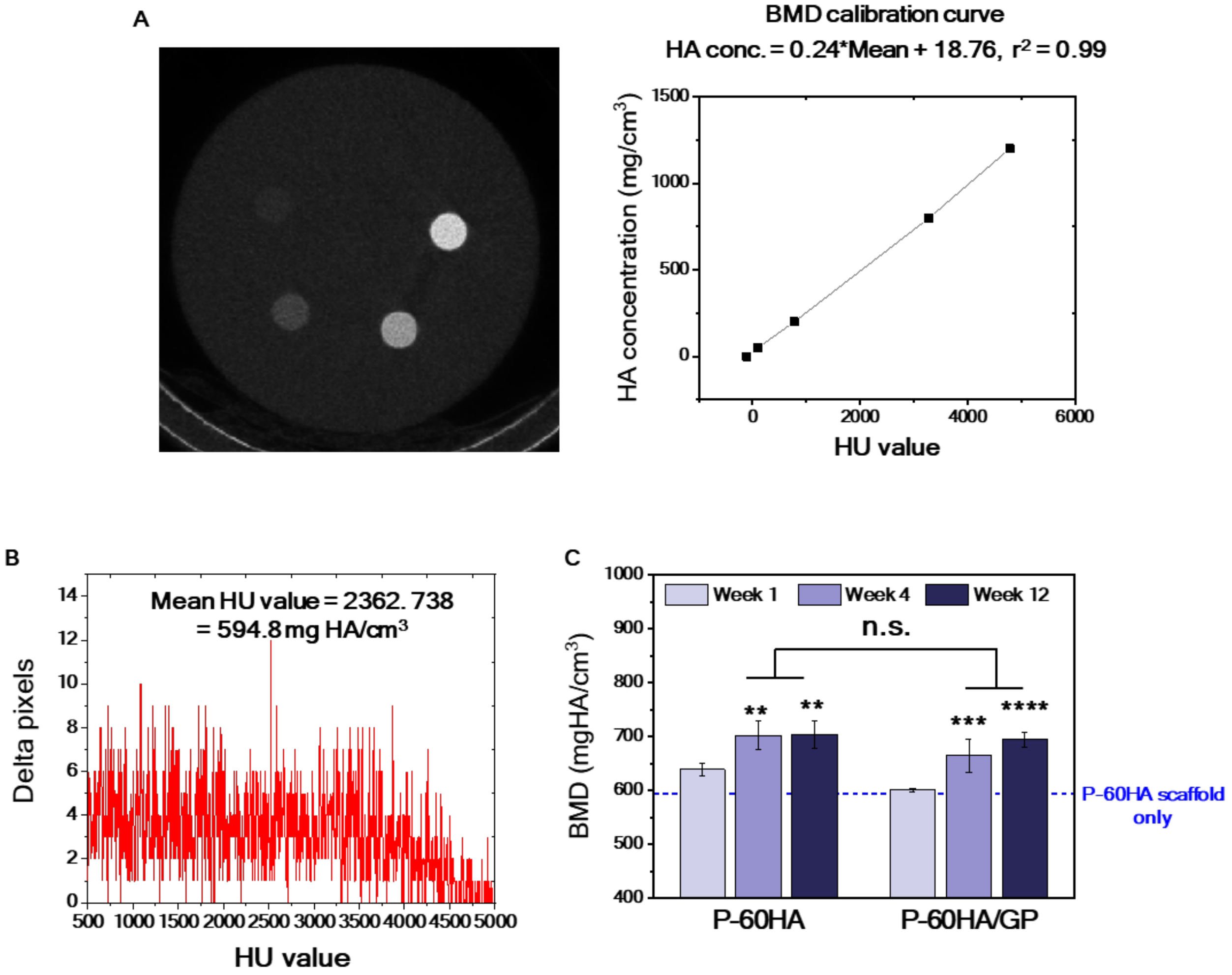


**Fig. S7. Conversion of Hounsfield unit (HU) values into bone mineral density (mg HA/cm$^3$).** (**A**) Representative images from phantom scanning and its scanned HU values converted into units of bone mineral density. (**B**) µCT scanning data plotted as delta pixels versus HU values for the P-60HA scaffold only. By using the calibration equation, the mean HU value is converted to BMD in mg HA/cm$^3$. (**C**) BMD measurements of newly formed tissue including scaffold for groups with P-60HA scaffolds. $^{**}p$ <0.01, $^{***}p$ <0.001 and $^{****}p$ <0.0001; n.s.: no significant difference; Error bars, ± SD; n=5.

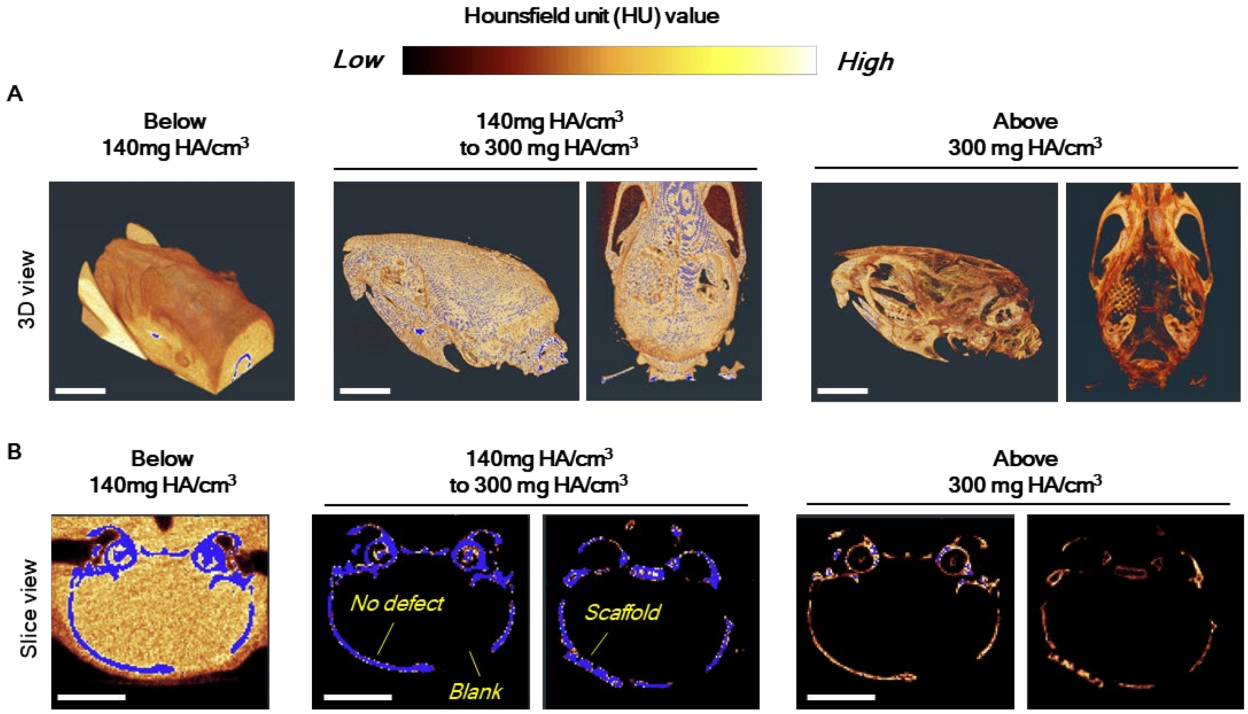


**Fig. S8. Visualized tissue area according to HU value of micro-computed tomography (μCT) scanning.** (**A**) 3D reconstructed images represented in regions of different HU values, and (**B**) 2D ortho-slice images depicting each range. Tissues are represented in a heat color map within the selected area, while unselected tissue areas are displayed in blue across the image. Scale bars, 5mm.

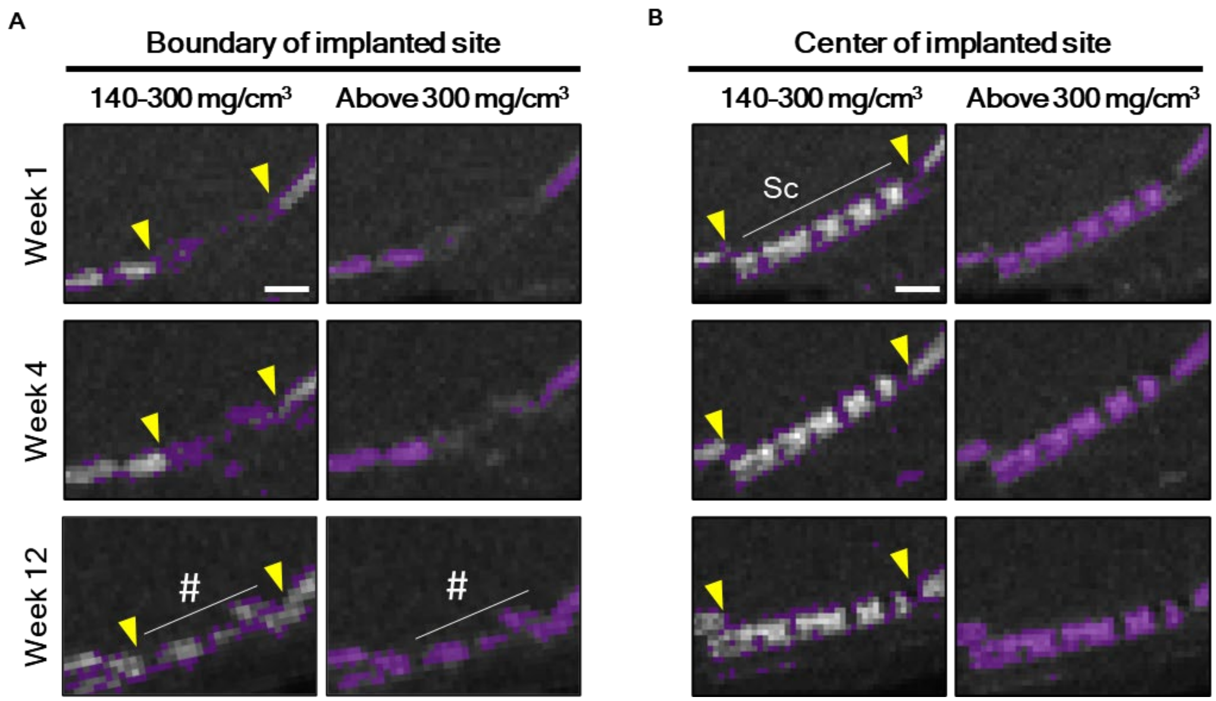


**Fig. S9. Visualized tissue area of the P-60HA/GP group over time at different HU values scanned by micro-computed tomography (μCT).** (**A**) Tissue morphology at the boundary and (**B**) the center of the implanted site. Corresponding tissue within the threshold range are represented in purple, while unselected tissue regions are displayed in grayscale. The comparison was performed from the same animal at the same location. Yello arrow: defect area, Hash (#): matured low-density bone, Sc: scaffold. Scale bars, 1mm.

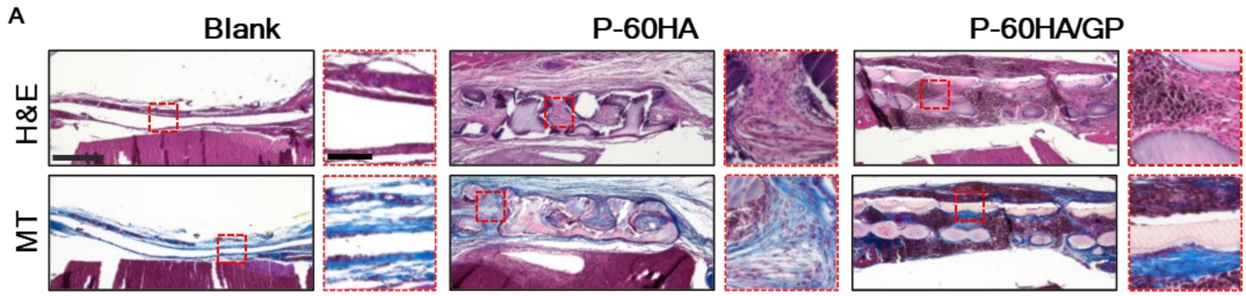


**Fig. S10. H&E and Masson's trichrome staining of Blank, P-60HA, and P-60HA/GP group at 4-week time point.** (A) Cross-sectional histological images of scaffold implanted tissue at 4 weeks. The images at the top and bottom represent H&E and Masson's trichrome staining images of each group, respectively. Scale bars, 500μm and 100μm (high magnification).

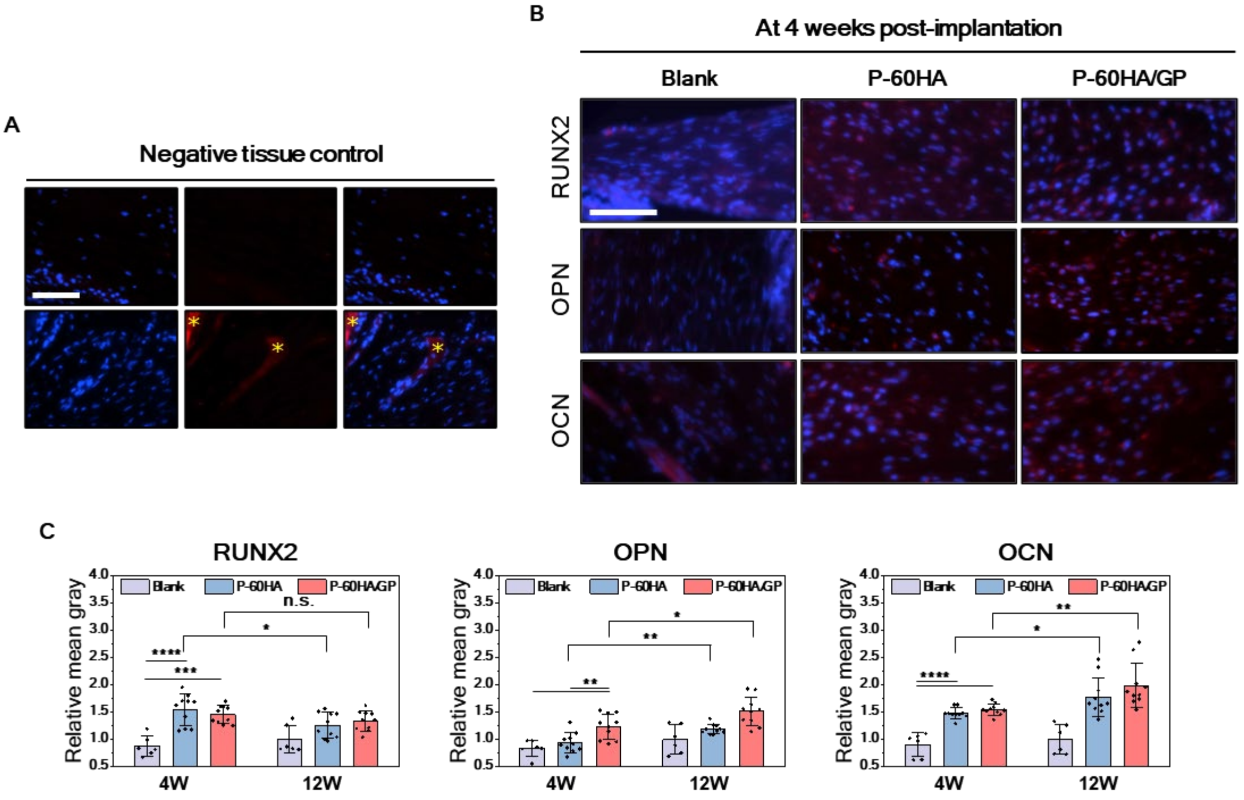


**Fig. S11. Immunofluorescence staining of Blank, P-60HA, and P-60HA/GP group at 4-week time point.** (**A**) Negative control of immunofluorescence. The asterisk (*): scaffold. Scale bar, 100μm. (**B**) Representative images of each osteogenic marker at 4 weeks. Scale bar, 100μm. (**C**) Quantification results of different markers for each group. $^{*}p$ <0.05, $^{**}p$ <0.01, $^{***}p$ <0.001 and $^{****}p$ <0.0001; n.s.: no significant difference; Error bars, ± SD; n=10.

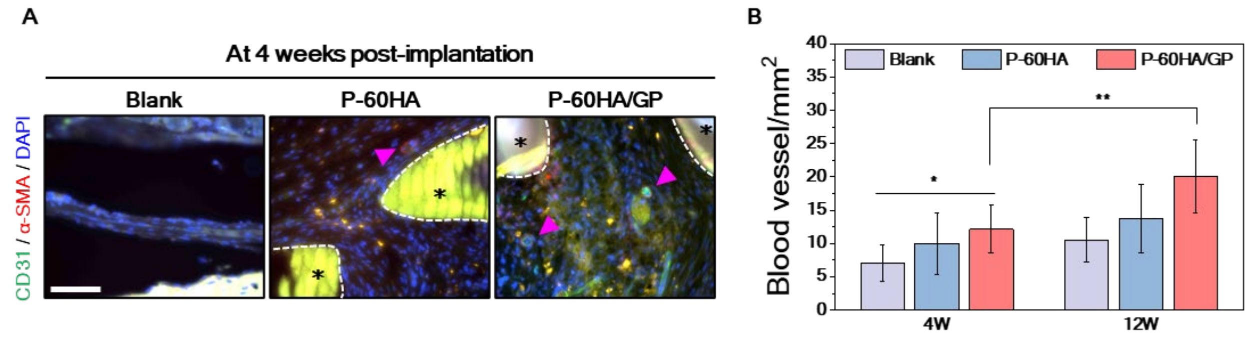


**Fig. S12. The assessment of blood vessel formation at 4-week time point.** (**A**) Representative images of each group at 4 weeks. The asterisk (*): scaffold; Pink arrow: blood vessel. Scale bar, 100µm. (**B**) Quantification results of blood vessel for each group. $^{*}p$ <0.05, $^{**}p$ <0.01; Error bars, ± SD; n=10.

| Gene | Forward primer sequence | Reverse primer sequence |
|---|---|---|
| Runt-related transcription factor2 (RUNX2) | GGTCAGATGCAGGCGGCCC | TACGTGTGGTAGCGCGTGGC |
| Osteopontin (OPN) | ACCTGAACGCGCCTTCTG | CATCCAGCTGACTCGTTTCATAA |
| Osteocalcin (OCN) | AGCAAAGGTGCAGCCTTTGT | GCGCCTGGGTCTCTTCACT |
| GAPDH (housekeeping) | AAGGTGAAGGTCGGAGTCAA | AATGAAGGGGTCATTGATGG |

**Table S1. Osteogenic gene sequences for RT-qPCR used to assess in vitro osteogenic differentiation of hMSCs.**
